# Regulator of G Protein Signaling 14 protein expression profile in the adult mouse brain

**DOI:** 10.1101/2024.06.22.600169

**Authors:** Sara N. Bramlett, Shana M. Fitzmaurice, Nicholas H. Harbin, Wuji Yan, Charan Bandlamudi, G. Emme Van Doorn, Yoland Smith, John R. Hepler

## Abstract

Regulator of G protein signaling 14 (RGS14) is a multifunctional signaling protein that serves as a natural suppressor of synaptic plasticity in the mouse brain. Our previous studies showed that RGS14 is highly expressed in postsynaptic dendrites and spines of pyramidal neurons in hippocampal area CA2 of the developing mouse brain. However, our more recent work with adult rhesus macaque brain shows that RGS14 is found in multiple neuron populations throughout hippocampal area CA1 and CA2, caudate nucleus, putamen, globus pallidus, substantia nigra, and amygdala in the adult rhesus monkey brain. In the mouse brain, we also have observed RGS14 protein in discrete limbic regions linked to reward behavior and addiction, including the central amygdala and the nucleus accumbens, but a comprehensive mapping of RGS14 protein expression in the adult mouse brain is lacking. Here, we report that RGS14 is more broadly expressed in mouse brain than previously known. Intense RGS14 staining is observed in specific neuron populations of the hippocampal formation, amygdala, septum, bed nucleus of stria terminalis and ventral striatum/nucleus accumbens. RGS14 is also observed in axon fiber tracts including the dorsal fornix, fimbria, stria terminalis, and the ventrohippocampal commissure. Moderate RGS14 staining is observed in various other adjacent regions not previously reported. These findings show that RGS14 is expressed in brain regions that govern aspects of core cognitive functions such as sensory perception, emotion, memory, motivation, and execution of actions, and suggests that RGS14 may serve to suppress plasticity and filter inputs in these brain regions to set the overall tone on experience-to-action processes.

## 1. INTRODUCTION

The Regulators of G protein Signaling (RGS) family of proteins serve to negatively regulate receptor and G protein signaling events (Hollinger et al., 2001; Sjögren, 2011; Stewart & Fisher, 2015). Their eponymous RGS domain facilitates GTP hydrolysis by active Gα-GTP subunits, greatly accelerating conversion to the inactive GDP-bound state. While many RGS proteins are dedicated GTPase activating proteins consisting of a single RGS domain, some are much larger and more complex featuring multiple domains and binding partners. One such RGS protein is RGS14, a multifunctional scaffolding protein enriched in the brain (Hollinger et al., 2001).

Within the brain, RGS14 is highly expressed in area CA2 of the hippocampus. Although RGS14 protein is expressed elsewhere outside of the central nervous system (Agudelo et al., 2018; Friedman et al., 2022; Li et al., 2016; Zhang et al., 2022), RGS14 actions are best understood in the brain where it integrates G protein, H-Ras/extracellular signal-regulated kinase (ERK) and Ca^2+^/calmodulin (CaM) signaling pathways to negatively regulate postsynaptic signaling and plasticity in neurons (Evans et al., 2018a; Harbin et al., 2021; Shu et al., 2010; Vellano et al., 2011; Vellano et al., 2013). RGS14 is highly enriched in dendritic spines of hippocampal area CA2 where it acts to suppress long-term potentiation and structural plasticity linked to spatial learning (Evans et al., 2018b; Lee et al., 2010).

We previously reported the expression profile of RGS14 protein in the developing mouse brain, showing strong expression in hippocampal area CA2 and primary olfactory areas, piriform cortex, and entorhinal cortices (Evans et al., 2014). More recently, we reported that RGS14 is also expressed in discrete limbic structures in the adult mouse brain where it modulates locomotor behavior linked to environmental novelty and cocaine (Foster et al., 2021). However, the protein distribution pattern of RGS14 throughout the adult mouse brain is not well characterized. To fill this knowledge gap, the present study employs immunostaining and fluorescence microscopy using tissue from both wild-type (WT) and L507R (LR) mutant mice to provide a comprehensive map of the RGS14 protein expression profile in the adult mouse brain. RGS14 is a cytoplasmic-nuclear shuttling protein (Branch & Hepler, 2017; Cho et al., 2005), and the LR mutant features a CRISPR-generated alanine-to-arginine point mutation in the *rgs14* gene that disrupts nuclear export, thus trapping RGS14 protein inside the nucleus of its originating cell (Squires et al., 2020). Use of this RGS14-reporter mouse line benefits our approach in several ways: (1) it allows for the precise localization of RGS14-producing neuron populations, (2) the subcellular nuclear concentration of RGS14 in LR mice allows for detection in regions with low protein expression, and (3) the nuclear sequestration of RGS14 in LR mice permits their use as an additional negative control when comparing with WT immunoreactivity.

Using these tools and approaches, we show that the staining pattern of RGS14 protein in the adult mouse brain is consistent with that previously reported in monkey and human brains (Squires et al., 2018), but with notable staining in some brain regions not explored in these studies. The revised expression profile described herein implicates RGS14 with previously unknown central nervous system functions.

## 2. MATERIALS AND METHODS

### Animals and tissue collection

All animal housing and procedures were designed according to the National Institutes of Health Guidelines for the Care and Use of Laboratory Animals and were approved by the Emory University Institutional Animal Care and Use Committee. L507R (LR) mice were maintained on a C57BL/6J background and socially housed with wild-type (WT) littermates.

Tissue was harvested from 10 mice in total: 4 WT and 4 LR for RGS14 detection, and 2 WT for negative controls. All mice were adults (aged >3 months) and all groups were split evenly between males and females. Mice were intraperitoneally injected with 200 mg/kg sodium pentobarbital and transcardially perfused with 4% paraformaldehyde (PFA) in 0.01 M phosphate-buffered saline (PBS). Brains were extracted and post-fixed by submersion in the PFA solution for 24 hours, then sunk in 15% sucrose/PBS for 24 hours, then 30% sucrose/PBS for ≥24 hours (all at 4°C). Brains were embedded in Optimal Cutting Temperature compound (Tissue-Tek, Torrance, CA) on dry ice, then sectioned in the coronal plane at 40 µm with a cryostat (−20°C). Tissue sections were stored in PBS+0.025% sodium azide prior to staining.

### Immunohistochemistry

Tissue sections for RGS14 detection were submersed in 10 mM sodium citrate buffer for the purpose of antigen retrieval (3 min at 100°C). They were then washed in PBS and blocked for 1 hour in 5% normal goat serum (NGS)/PBS (room temp), then incubated in mouse α-RGS14 primary antibody (NeuroMab, Davis, CA, clone N133/21, RRID AB_2877352; 1:500) in NGS/PBS at 4°C overnight. This monoclonal antibody was previously characterized in great depth and shown to be highly specific for rodent RGS14 (Evans et al., 2014). The following day, the sections were washed in PBS and incubated in goat α-mouse AlexaFluor 488+ secondary antibody (Invitrogen, 1:500) in NGS/PBS for 2 hours at room temp, washed again in PBS, and mounted onto Superfrost Plus slides (Thermo Fisher Scientific, Waltham, MA). Once dry, slides were coverslipped with Fluoromount-G+DAPI mounting medium (Southern Biotech, Birmingham, AL) and stored at −4°C until imaging. An identical protocol was followed for negative control sections, excluding overnight incubation in primary antibody.

### Fluorescent imaging and image processing

Sections were imaged using two microscopes on separate occasions. Initially, they were imaged with a Leica DM6000 epifluorescent upright microscope at 5x and 10x magnification, using identical exposure settings between subjects for each anatomical region. Later, they were imaged with a Leica SP8 MP multiphoton confocal microscope using the tile function with automatic z-axis adjustment at each position to stitch together adjacent 10x magnified images (smooth stitching at 10% overlap). Exposure and image acquisition settings for heat atlas images were kept largely consistent. Certain sources of variability during heat atlas image acquisition were unavoidable, but were individually assessed and corrected for when possible. Their impact overall was determined to be minor if not negligible. Examples of such sources include between-session instrument variability, variation in autofluorecence intensity, and slight photobleaching effects in tissue imaged repeatedly.

Microscopy images were post-processed using ImageJ and figures were composed using Inkscape. All images were compared to negative controls (lacking primary antibody) to identify nonspecific staining. Apart from slight corrections for the aforementioned minor sources of variability, pixel value ranges were kept consistent between heat atlas images to allow for relative quantification of RGS14 levels throughout the brain from pseudo-standardized images. Pixel value ranges for non-heat atlas images were set relative to each image in order to maximize signal-to-noise, while still maintaining the appearance of identifying structural features. Heat atlas images were derived from a single WT female subject whose tissue had been sectioned at an optimal angle for alignment with the coronal Allen Mouse Brain Atlas. RGS14 expression in this mouse reflected that of the other WT subjects, and no sex differences were seen. Non-heat atlas images were obtained from a variety of subjects, both male and female.

The ImageJ pipeline for heat atlas images included: despeckle → gaussian blur filter → adjust brightness/contrast → apply heat map look-up table (LUT). The processed image was then overlaid onto the grayscale original image. The heat map LUT used is the Jet LUT native to ImageJ with the low end extended into black, which allows for transparency when overlaid. Heat atlas images were juxtaposed with matching Allen Mouse Brain Atlas graphics in Inkscape. Allen Brain Atlas graphics were manually warped using lattice deformation to align with heat atlas images. The ImageJ pipeline for non-heat atlas images included: despeckle → median filter → remove outliers (for artifact elimination) → adjust brightness/contrast → apply Virdis LUT. Occasionally, background was subtracted using a rolling ball filter in LR tissue images to reduce autofluorescence background. Image post-processing was performed with the explicit goal of providing data clarity while maintaining data integrity. Raw files of all images are available upon request.

All images were matched to the Allen Mouse Brain Atlas. Regions exhibiting known or suspected false positive signal based on negative control staining were excluded from the report. Naming conventions largely followed the Allen Mouse Brain Atlas, with a few minor exceptions.

### Assessment of relative protein levels

Hemispheric images were acquired and pseudo-standardized as described above. The heat map LUT was applied to each image, and RGS14 level as determined by relative fluorescence intensity was assessed for each brain region. Relative protein was binned into five incremental levels, listed here in ranking order from lowest to highest protein levels by the terminology used to describe them in the text and the corresponding color in heat atlas and tables: light (dark blue), low (teal), moderate (green), high (yellow), and intense (red). For simplicity of the tabulated results, regions with variable RGS14 levels were binned based on overall average level. Heat atlas images presented in the figures include a representational selection of those used to assess relative protein levels.

Within each subsection of the results main text, ordinal organization of regions described generally follows the format of descending order of RGS14 level, followed by position where the region first appears along the rostro-caudal axis. Tabulated results are also initially ordered by descending RGS14 level, but follow an alphabetical second order of organization for ease of reference. Imaging sub-figures are ordered rostrocaudally. Tables were composed using the ggtable and fontawesome packages for R.

## 3. RESULTS

### Heat atlas of RGS14 protein

Figures 1 & 2 depict a series of RGS14-immunostained coronal sections through the rostro-caudal axis of the mouse brain up to the anterior region of the cerebellum. This comprehensive atlas of RGS14 protein levels reveals that RGS14 is more widely expressed in the adult mouse brain than previously known. Substantial RGS14 immunoreactivity is seen in regions of allocortex, septum, striatum, pallidum, and to a lesser extent ventral midbrain. Widespread, but lighter, immunoreactivity is seen across isocortex. The protein is also found at varying degrees along many of the axonal fiber tracts that run between these regions. In terms of functional anatomy, RGS14 is largely found in regions of limbic system, basal ganglia, and sensory, motor, and association cortices. The relative intensity of RGS14 immunoreactivity across all examined regions is summarized in Tables 1 & 2. The cellular localization of RGS14 labeling in specific brain regions is discussed in the following sections.

**Fig. 1:**
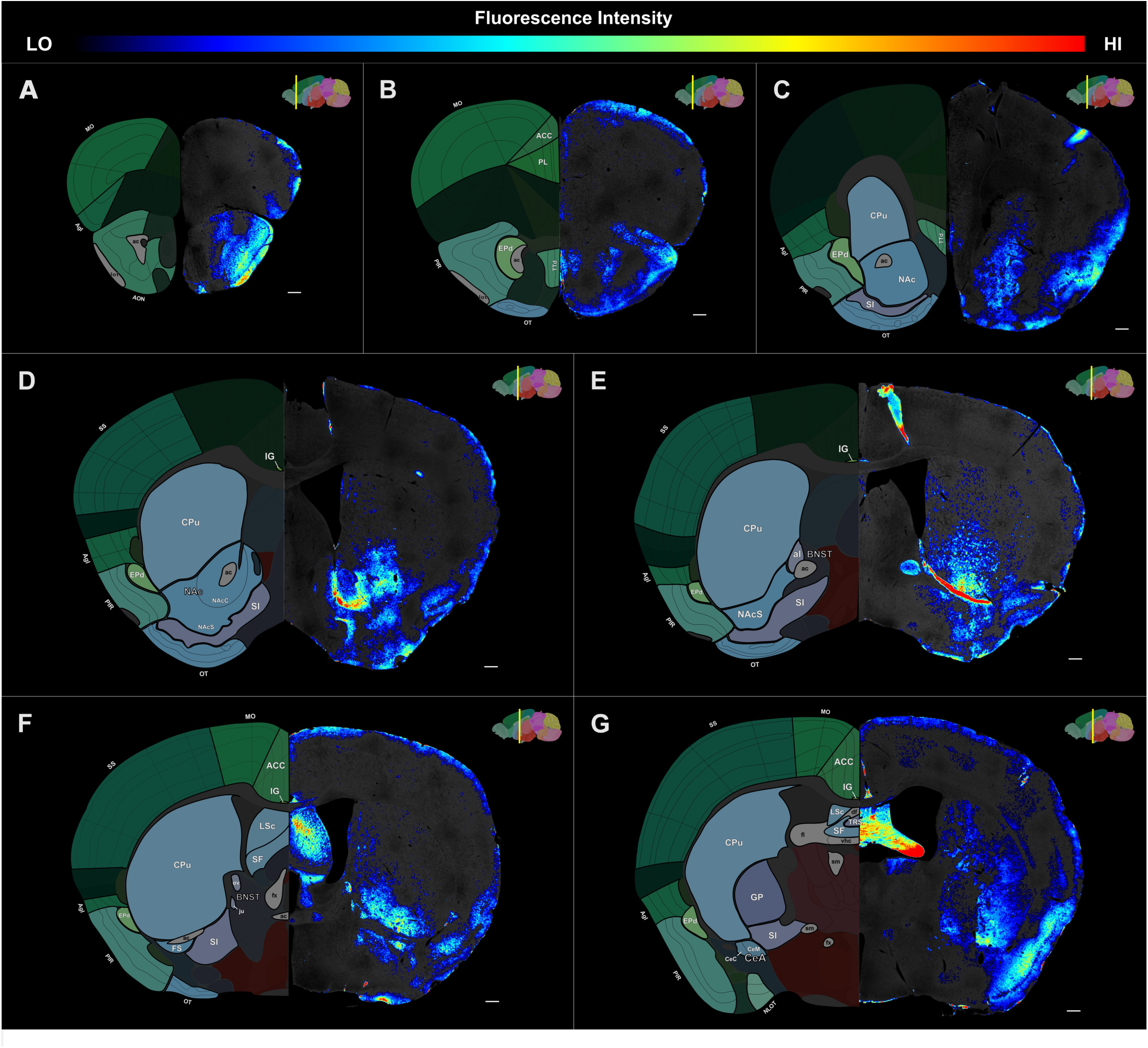
Heat atlas of RGS14 protein levels in adult mouse brain: rostral to medial. Relative fluorescence intensity in RGS14-immunostained coronal brain sections from a wild type (WT) mouse in series across the rostro-caudal axis (**A** → **G**). RGS14 (right) and anatomical annotations (left) from the Allen Mouse Brain Reference Atlas (atlas.brain-map.org). RGS14 immunoreactivity is seen within the confines of the regions labeled on the reference atlas. Notable immunoreactivity is seen in olfactory cortex, striatum, and septum, with lighter staining in isocortex. Light text indicates nuclei and subnuclei; dark text indicates fiber tracts. Yellow bar indicates sagittal position. Scale bar = 100 µm. Abbreviations: <u>regions:</u> **ACC**: anterior cingulate cortex, **AgI**: agranular insular cortex, **AON**: anterior olfactory nucleus, **BNST**: bed nucleus of the stria terminals, **CeA**: central amygdala, **Ce(C,M)**: central amygdala- (capsular, medial), **CPu**: caudoputamen, **EPd**: endopiriform nucleus-dorsal, **FS**: fundus of the striatum, **GP**: globus pallidus, **IG**: induseum griseum, **ju**: juxtacapsular nucleus of BNST, **LSc**: lateral septal nucleus-caudal, **MO**: motor cortex, **NAc(C,S)**: nucleus accumbens- (core, shell), **NLOT**: nucleus of the lateral olfactory tract, **OT**: olfactory tubercle, **ov**: oval nucleus of the BNST, **PIR**: piriform cortex, **PL**: prelimbic cortex, **SF**: septofimbrial nucleus, **SI**: substantia innominata, **SS**: somatosensory cortex, **TRS**: triangular nucleus of the septum, **TTd**: taenia tecta-dorsal; <u>fiber tracts:</u> **ac**: anterior commissure, **df**: dorsal fornix, **fi**: fimbria, **fx**: fornix, **lot**: lateral olfactory tract, **sm**: stria medullaris, **vhc**: ventral hippocampal commissure.

**Fig. 2:**
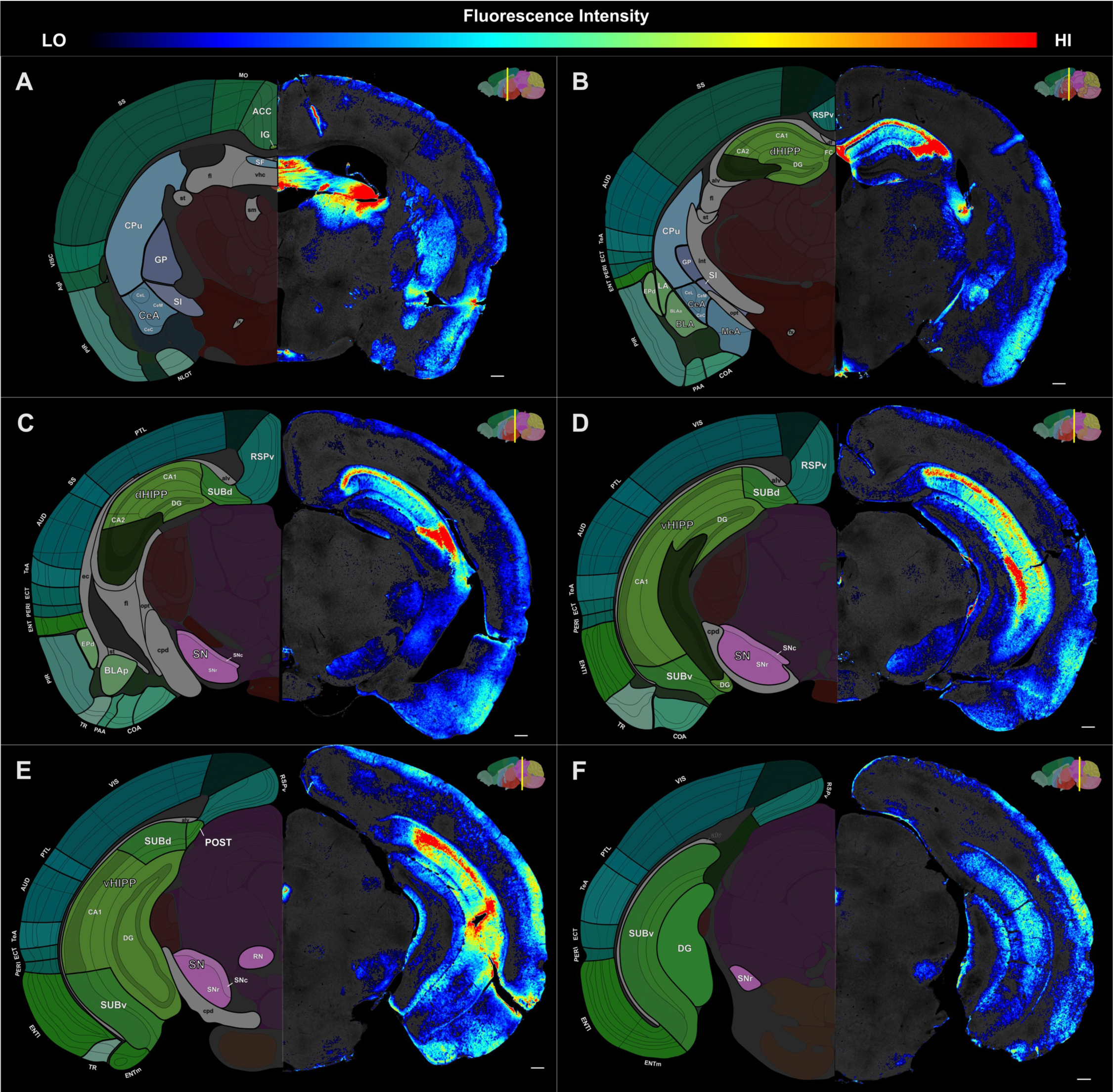
Heat atlas of RGS14 protein levels in adult mouse brain: medial to caudal. Relative fluorescence intensity in RGS14-immunostained coronal brain sections from a wild type (WT) mouse in series across the rostro-caudal axis (**A** → **F**). RGS14 (right) and anatomical annotations (left) from the Allen Mouse Brain Reference Atlas (atlas.brain-map.org). RGS14 immunoreactivity is seen within the confines of the regions labeled on the reference atlas. Notable RGS14 immunoreactivity is seen in septum, hippocampal formation, and amygdala, with lighter staining in isocortex and discrete midbrain subregions. Light text indicates nuclei and subnuclei; dark text indicates fiber tracts. Yellow bar indicates sagittal position. Scale bar = 100 µm. Abbreviations: <u>regions:</u> **ACC**: anterior cingulate cortex, **AUD**: auditory cortex, **AgI**: agranular insular cortex, **BLA(a,p)**: basolateral amygdala- (anterior, posterior), **CA(1,2)**: hippocampal field *cornu ammonis* (1, 2), **CeA**: central amygdala, **Ce(C,L,M)**: central amygdala- (capsular, lateral, medial), **COA**: cortical amygdalar area, **CPu**: caudoputamen, **DG**: dentate gyrus, **dHIPP**: dorsal hippocampus, **ECT**: ectorhinal cortex, **ENT(l,m)**: entorhinal cortex- (lateral, medial), **EPd**: endopiriform nucleus-dorsal, **FC**: fasciola cinerea, **GP**: globus pallidus, **IG**: induseum griseum, **LA**: lateral amygdala, **MeA**: medial amygdala, **MO**: motor cortex, **NLOT**: nucleus of the lateral olfactory tract, **PAA**: piriform-amygdalar area, **PERI**: perirhinal cortex, **PIR**: piriform cortex, **POST**: postsubiculum, **PTL**: posterior parietal association cortex, **RN**: red nucleus, **RSPv**: retrosplenial cortex-ventral, **SF**: septofimbrial nucleus, **SI**: substantia innominata, **SN(c,r)**: substantia nigra- (pars compacta, pars reticulata), **SS**: somatosensory cortex, **SUB(d,v)**: subiculum- (dorsal, ventral), **TeA**: temporal association cortex, **TR**: postpiriform transition cortex, **vHIPP**: ventral hippocampus, **VIS**: visual cortex, **VISC**: visceral cortex; <u>fiber tracts:</u> **alv**: alveus, **cpd**: cerebral peduncle, **df**: dorsal fornix, **ec**: external capsule, **fi**: fimbria, **fx**: fornix, **int**: internal capsule, **opt**: optic tract, **sm**: stria medullaris, **st**: stria terminalis, **vhc**: ventral hippocampal commissure.

**Table 1:**
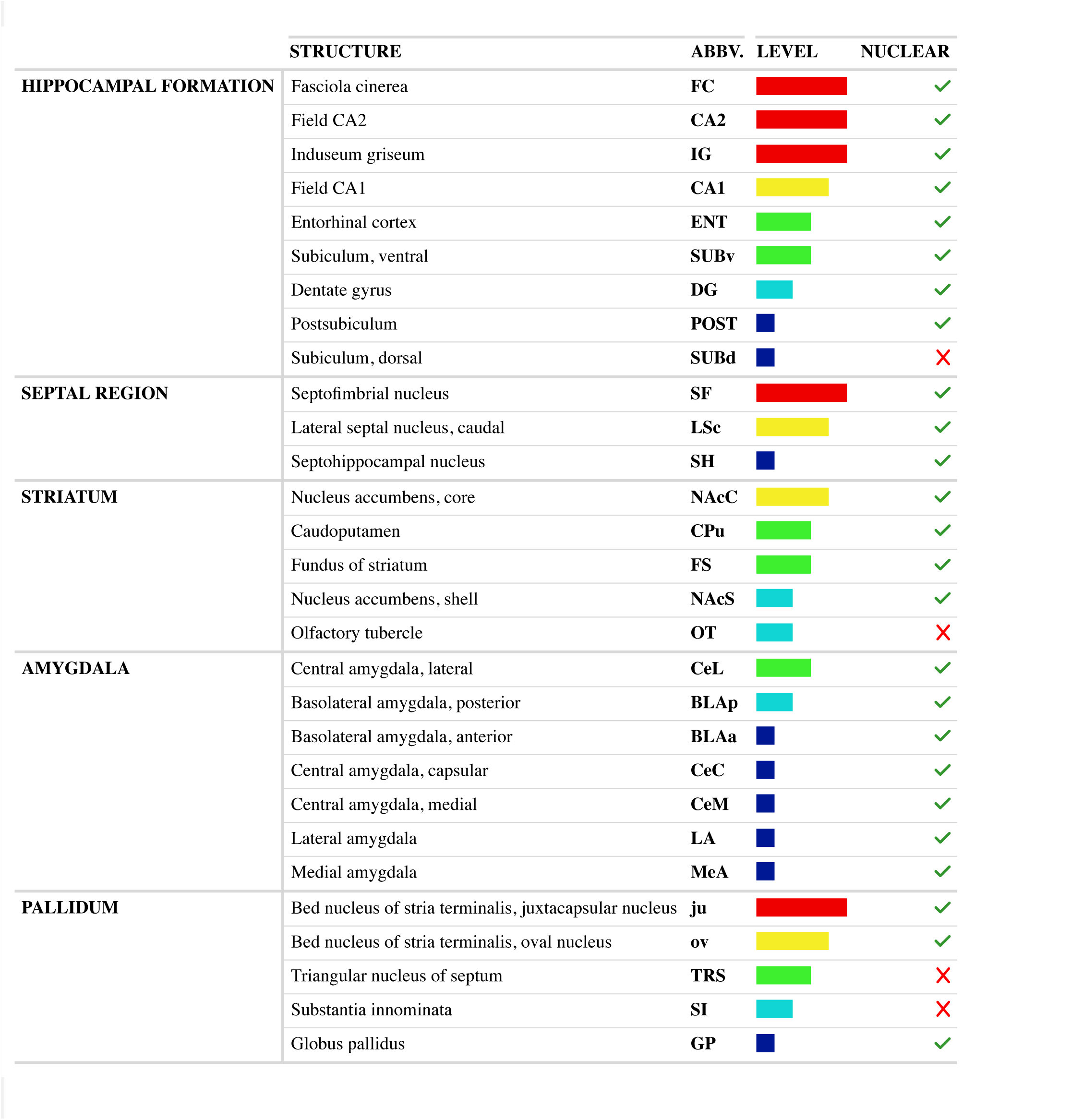
Summary of RGS14 expression in adult mouse brain: hippocampus, septum, striatum, amygdala, and pallidum. Relative expression determined from immunofluorescence heat atlas.

|  | STRUCTURE | ABBV. | LEVEL | NUCLEAR |
| --- | --- | --- | --- | --- |
| HIPPOCAMPAL FORMATION | Fasciola cinerea | FC |  | ✓ |
|  | Field CA2 | CA2 |  | ✓ |
|  | Induseum griseum | IG |  | ✓ |
|  | Field CA1 | CA1 |  | ✓ |
|  | Entorhinal cortex | ENT |  | ✓ |
|  | Subiculum, ventral | SUBv |  | ✓ |
|  | Dentate gyrus | DG |  | ✓ |
|  | Postsubiculum | POST |  | ✓ |
|  | Subiculum, dorsal | SUBd |  | ✗ |
| SEPTAL REGION | Septofimbrial nucleus | SF |  | ✓ |
|  | Lateral septal nucleus, caudal | LSc |  | ✓ |
|  | Septohippocampal nucleus | SH |  | ✓ |
| STRIATUM | Nucleus accumbens, core | NAcC |  | ✓ |
|  | Caudoputamen | CPu |  | ✓ |
|  | Fundus of striatum | FS |  | ✓ |
|  | Nucleus accumbens, shell | NAcS |  | ✓ |
|  | Olfactory tubercle | OT |  | ✗ |
| AMYGDALA | Central amygdala, lateral | CeL |  | ✓ |
|  | Basolateral amygdala, posterior | BLAp |  | ✓ |
|  | Basolateral amygdala, anterior | BLAa |  | ✓ |
|  | Central amygdala, capsular | CeC |  | ✓ |
|  | Central amygdala, medial | CeM |  | ✓ |
|  | Lateral amygdala | LA |  | ✓ |
|  | Medial amygdala | MeA |  | ✓ |
| PALLIDUM | Bed nucleus of stria terminalis, juxtacapsular nucleus | ju |  | ✓ |
|  | Bed nucleus of stria terminalis, oval nucleus | ov |  | ✓ |
|  | Triangular nucleus of septum | TRS |  | ✗ |
|  | Substantia innominata | SI |  | ✗ |
|  | Globus pallidus | GP |  | ✓ |

**Table 2:**
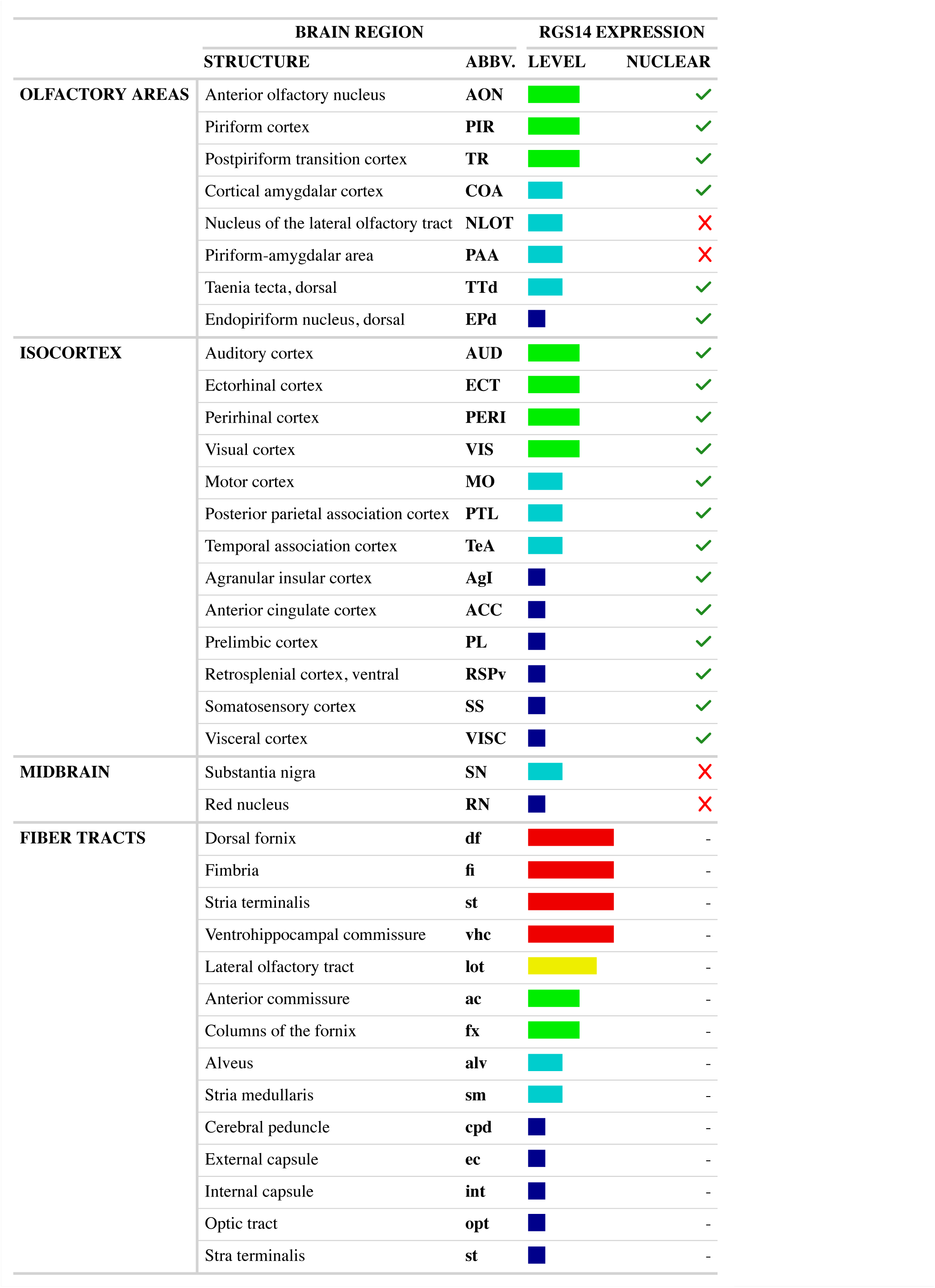
Summary of mouse brain RGS14 expression: olfactory area, isocortex, midbrain, and fiber tracts. Relative expression determined from immunofluorescence heat atlas.

|  | BRAIN REGION |  | RGS14 EXPRESSION |  |
| --- | --- | --- | --- | --- |
|  | STRUCTURE | ABBV. | LEVEL | NUCLEAR |
| OLFACTORY AREAS | Anterior olfactory nucleus | AON |  | ✓ |
|  | Piriform cortex | PIR |  | ✓ |
|  | Postpiriform transition cortex | TR |  | ✓ |
|  | Cortical amygdalar cortex | COA |  | ✓ |
|  | Nucleus of the lateral olfactory tract | NLOT |  | ✗ |
|  | Piriform-amygdalar area | PAA |  | ✗ |
|  | Taenia tecta, dorsal | TTd |  | ✓ |
|  | Endopiriform nucleus, dorsal | EPd |  | ✓ |
| ISOCORTEX | Auditory cortex | AUD |  | ✓ |
|  | Ectorhinal cortex | ECT |  | ✓ |
|  | Perirhinal cortex | PERI |  | ✓ |
|  | Visual cortex | VIS |  | ✓ |
|  | Motor cortex | MO |  | ✓ |
|  | Posterior parietal association cortex | PTL |  | ✓ |
|  | Temporal association cortex | TeA |  | ✓ |
|  | Agranular insular cortex | AgI |  | ✓ |
|  | Anterior cingulate cortex | ACC |  | ✓ |
|  | Prelimbic cortex | PL |  | ✓ |
|  | Retrosplenial cortex, ventral | RSPv |  | ✓ |
|  | Somatosensory cortex | SS |  | ✓ |
|  | Visceral cortex | VISC |  | ✓ |
| MIDBRAIN | Substantia nigra | SN |  | ✗ |
|  | Red nucleus | RN |  | ✗ |
| FIBER TRACTS | Dorsal fornix | df |  | - |
|  | Fimbria | fi |  | - |
|  | Stria terminalis | st |  | - |
|  | Ventrohippocampal commissure | vhc |  | - |
|  | Lateral olfactory tract | lot |  | - |
|  | Anterior commissure | ac |  | - |
|  | Columns of the fornix | fx |  | - |
|  | Alveus | alv |  | - |
|  | Stria medullaris | sm |  | - |
|  | Cerebral peduncle | cpd |  | - |
|  | External capsule | ec |  | - |
|  | Internal capsule | int |  | - |
|  | Optic tract | opt |  | - |
|  | Stria terminalis | st |  | - |

### Hippocampal formation

The intensity of RGS14 immunoreactivity in the hippocampal formation is generally high throughout, though the bulk of hippocampal formation RGS14 immunoreactivity is seen in the hippocampus proper. In the CA fields in WT animals, intense signal is found in CA2 (Fig. 3F), with lesser but still high signal in CA1, while CA3 is apparently devoid of labeling (Fig. 3F,I). This distribution pattern remains the same throughout the dorsoventral extent of the hippocampus. Signal is strongest in the pyramidal layer, then dissipates outwards through processes extending into the stratum oriens and stratum radiatum/stratum lacunosum moleculare. In line with this, examination of LR tissue reveals RGS14-positive nuclei in the pyramidal layer of CA2 (Fig. 3E) and CA1 (Fig. 3H & Fig. 6H). In WT mice, intense signal is additionally found within the fasciola cinerea (FC, Fig. 3C) and indusium griseum (IG, Fig. 5F & Fig. 10C). In LR mice, a dense layer of RGS14-positive nuclei is found in both regions (FC: Fig. 3B; IG: Fig. 5E & Fig. 10B). Diffuse low-level signal is seen in the polymorph and, particularly, the molecular layers of the dentate gyrus (DG) in WT animals (Fig. 3L), while RGS14-positive nuclei are predominantly found in the granule cell layer of LRs (Fig. 3K).

**Fig. 3:**
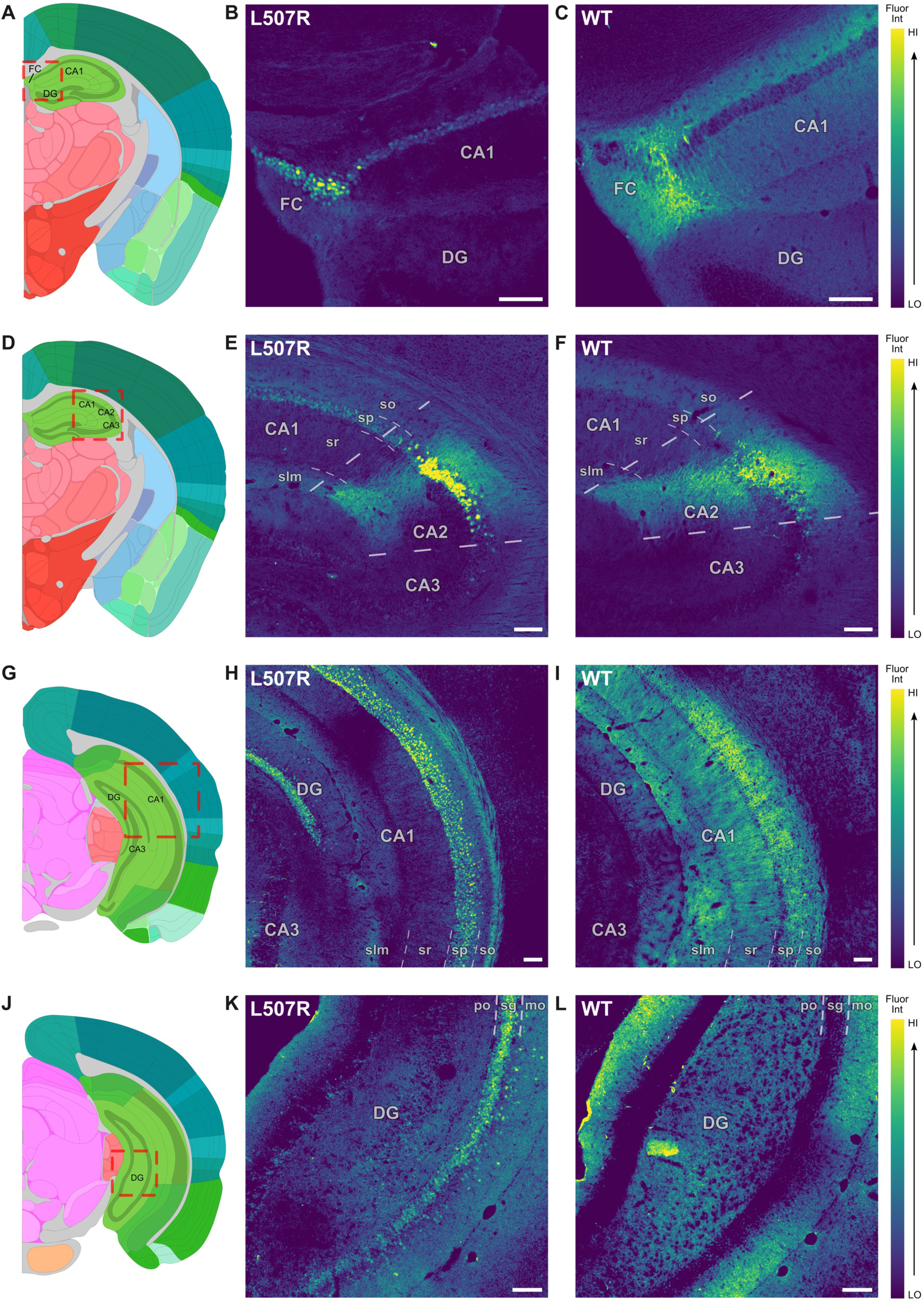
RGS14 is highly expressed in the bulk of hippocampus proper. Relative fluorescence intensity in RGS14-immunostained tissue sections from wild type (WT) and L507R-reporter mice. Anatomical annotations (left; **A,D,G,J**) from the Allen Mouse Brain Reference Atlas (atlas.brain-map.org) aligned with confocal images of RGS14 immunoreactivity in LR reporter (middle; **B,E,H,K**) and WT mice (right; **C,F,I,L**). Red dashed box in left panels indicates approximate anatomical location of tissue cross-sections in middle and right panels in each row. Scale bar = 100 µm. Abbreviations: **CA(1,2,3)**: hippocampal field *cornu ammonis* (1,2,3), **DG**: dentate gyrus, **FC**: fasciola cinerea; <u>layers:</u> **slm**: stratum lacunosum-moleculare, **sr**: stratum radiatum, **sp**: stratum pyramidale, **so**: stratum oriens; **po**: polymorph layer, **sg**: stratum granulosum, **mo**: molecular layer.

Overall moderate, but variable levels of RGS14 immunoreactivity are seen in retrohippocampal regions. Moderate signal is seen throughout the entorhinal cortex (ENT, Fig. 4I & Fig. 9K) in WT mice. Staining in LR tissue reveals RGS14-positive nuclei in most, if not all, ENT layers, with the exception of the most superficial layer I (Fig. 4H & Fig. 9J). Staining in WT subiculum is moderate in the ventral subregion (SUBv, Fig. 4F,I) but lighter in the dorsal subregion (SUBd, Fig. 2C & Fig. 3L). LR tissue reveals that RGS14-positive nuclei are seen exclusively in SUBv (Fig. 4E,Hd) and not in SUBd (Fig. 4B & Fig. 6E). A likely explanation is that RGS14 in SUBd is located within axonal and terminal processes from such SUBv and/or other projecting regions. Lastly, light staining is seen in the postsubiculum (POST, Fig. 4L) of WT mice, while RGS14-positive nuclei are dispersed across the region in LR animals (Fig. 4K).

**Fig. 4:**
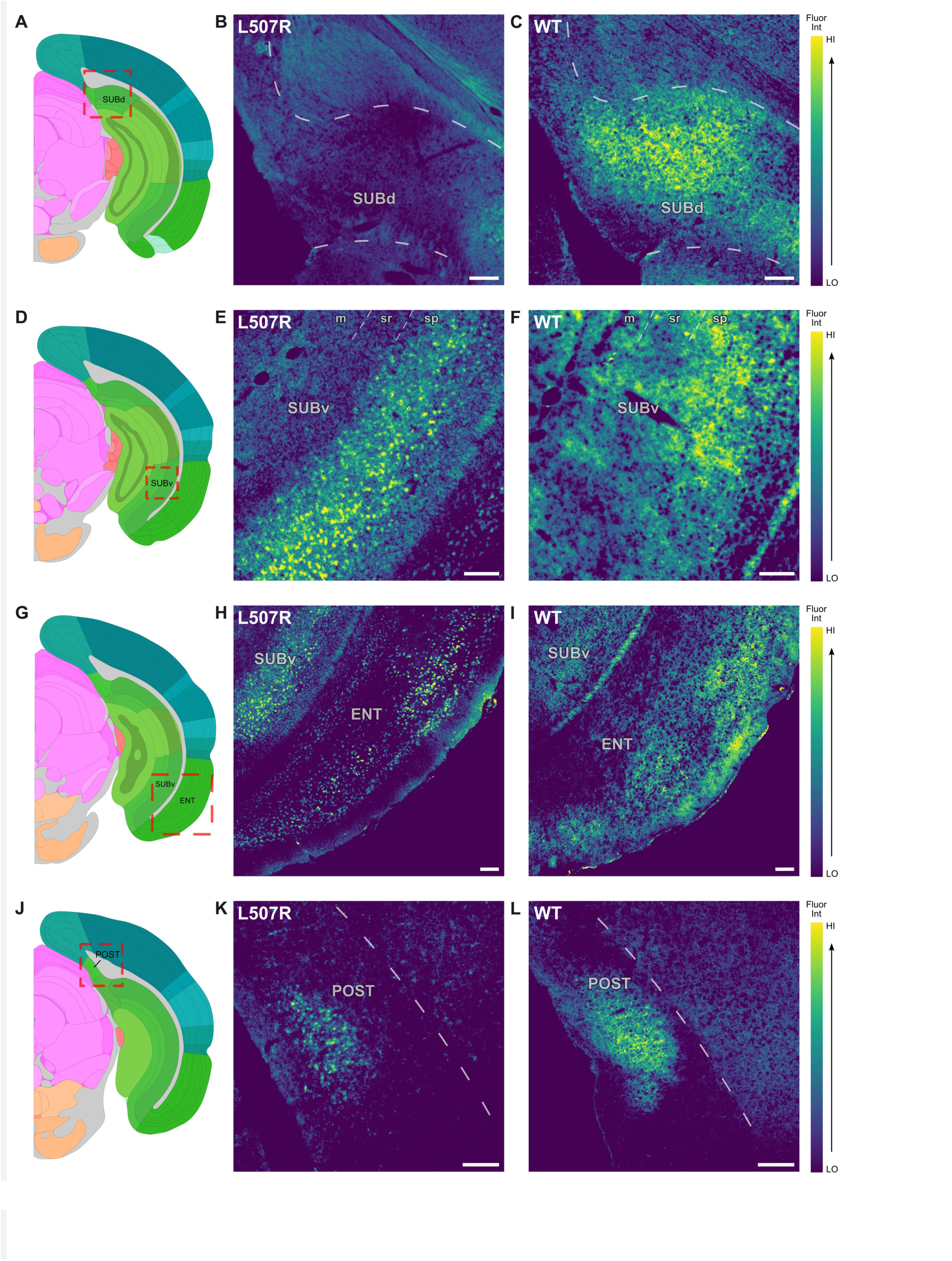
RGS14 is widely expressed at moderate levels in retrohippocampal regions. Relative fluorescence intensity in RGS14-immunostained tissue sections from wild type (WT) and L507R-reporter mice. Anatomical annotations (left; **A,D,G,J**) from the Allen Mouse Brain Reference Atlas (atlas.brain-map.org) aligned with confocal images of RGS14 immunoreactivity in LR reporter (middle; **B,E,H,K**) and WT mice (right; **C,F,I,L**). Red dashed box in left panels indicates approximate anatomical location of tissue cross-sections in middle and right panels in each row. Scale bar = 100 µm. Abbreviations: **ENT**: entorhinal cortex, **POST**: postsubiculum, **SUB(d,v)**: subiculum- (dorsal, ventral); <u>layers:</u> **m**: molecular layer, **sr**: stratum radiatum, **sp**: stratum pyramidale.

**Fig. 5:**
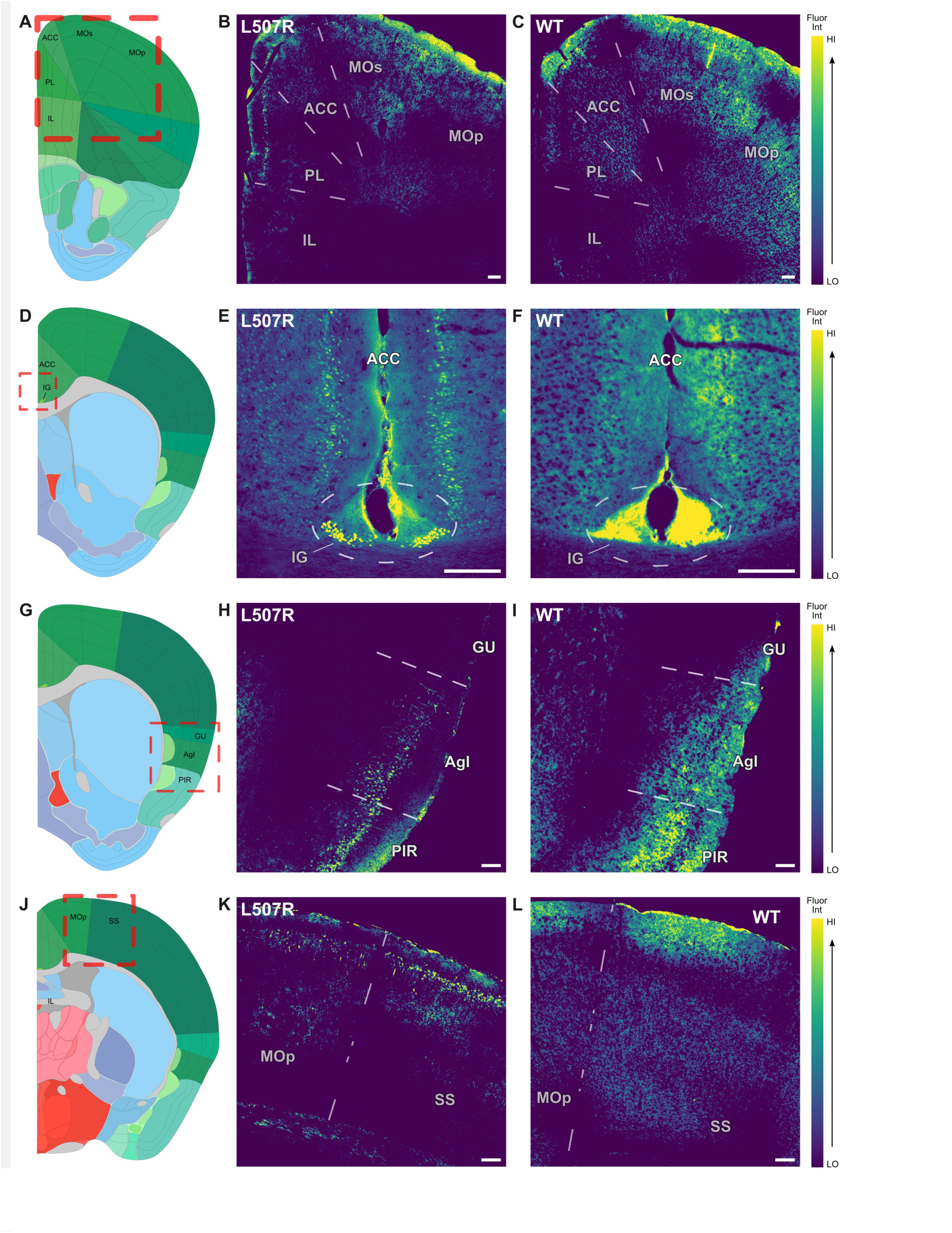
RGS14 is expressed at low levels in prefrontal cortex and frontal cortex. Relative fluorescence intensity in RGS14-immunostained tissue sections from wild type (WT) and L507R-reporter mice. Anatomical annotations (left; **A,D,G,J**) from the Allen Mouse Brain Reference Atlas (atlas.brain-map.org) aligned with confocal images of RGS14 immunoreactivity in LR reporter (middle; **B,E,H,K**) and WT mice (right; **C,F,I,L**). Red dashed box in left panels indicates approximate anatomical location of tissue cross-sections in middle and right panels in each row. Scale bar = 100 µm. Abbreviations: **ACC**: anterior cingulate cortex, **AgI**: agranular insular cortex, **GU**: gustatory cortex, **IG**: induseum griseum, **IL**: infralimbic cortex, **MO(p,s)**: motor cortex- (primary, secondary), **PIR**: piriform cortex, **PL**: prelimbic cortex, **SS**: somatosensory cortex.

**Fig. 6:**
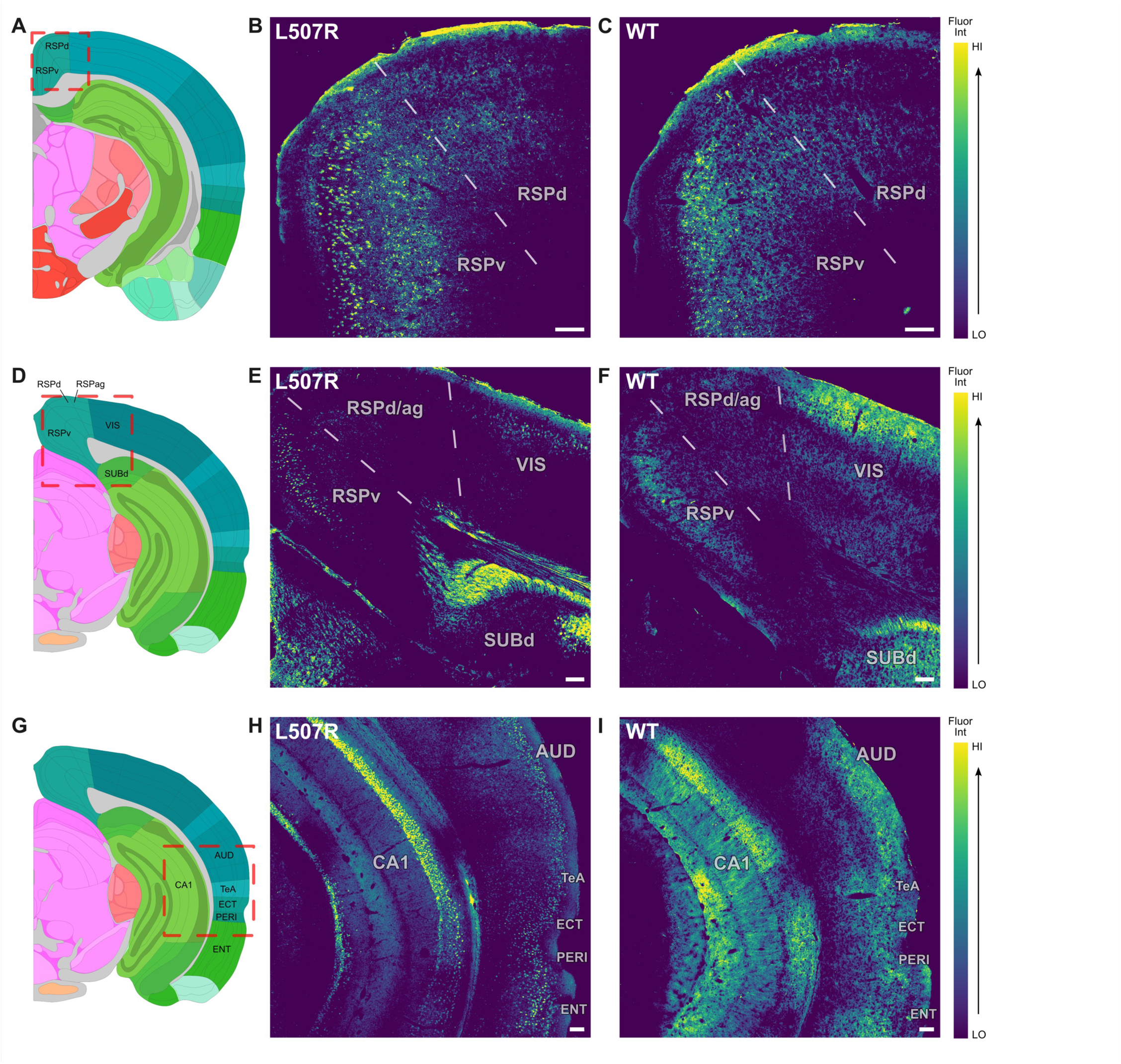
RGS14 is moderately expressed in parietal-temporal and occipital cortex. Relative fluorescence intensity in RGS14-immunostained tissue sections from wild type (WT) and L507R-reporter mice. Anatomical annotations (left; **A,D,G,J**) from the Allen Mouse Brain Reference Atlas (atlas.brain-map.org) aligned with confocal images of RGS14 immunoreactivity in LR reporter (middle; **B,E,H,K**) and WT mice (right; **C,F,I,L**). Red dashed box in left panels indicates approximate anatomical location of tissue cross-sections in middle and right panels in each row. Scale bar = 100 µm. Abbreviations: **AUD**: auditory cortex, **CA1**: hippocampal field *cornu ammonis* 1, **ECT**: ectorhinal cortex, **ENT**: entorhinal cortex, **PERI**: perirhinal cortex, **RSP(d,v,ag)**: retrosplenial cortex- (dorsal, ventral, agranular), **SUBd**: subiculum-dorsal, **TeA**: temporal association cortex, **VIS**: visual cortex.

### Isocortex

RGS14 immunoreactivity is broadly seen across the isocortex, though at much more modest levels than in allocortex. In WT mice, light signal is seen in the prefrontal cortex, specifically in the prelimbic (PL, Fig. 5C), anterior cingulate (ACC, Fig. 5C,F), and agranular insular (AgI, Fig. 5I) areas, though apparently absent in the infralimbic cortex (Fig. 5C). Signal is somewhat higher, though still low, in association areas beyond the frontal lobe, namely the temporal association (TeA) and posterior parietal association cortices (PTL, Fig. 6I). The strongest RGS14 labeling in association cortices is seen at moderate levels in the ectorhinal (ECT) and perirhinal (PERI, Fig. 6I) cortex. This is unsurprising given the close anatomo-functional relationship of these regions with the hippocampal formation. Light signal is also seen exclusively in the ventral subregion of the retrosplenial cortex (RSPv, Fig. 6C,F). RGS14 is seen in sensory cortical regions, including moderate levels in visual (VIS, Fig. 6F) and auditory (AUD, Fig. 6I) cortices and light levels in the somatosensory (SS, Fig. 5L) and visceral (VISC, not shown) cortex. It is also found at varying, but overall low, levels in primary and secondary motor cortices (MO) (Fig. 5C,L). We previously noted a similar protein distribution pattern across the isocortex in adolescent mice, but it was transient and undetectable in adults (Evans et al., 2014).

The data from the LR mice in the present study show that, while mostly subtle, RGS14 is unequivocally expressed in the adult mouse isocortex. Figure 6H particularly exemplifies the consistent pattern of the bulk of RGS14-positive nuclei residing in more superficial layers below layer I, presumably layer II/III, with fewer labeled nuclei in deeper layers. In WT mice, RGS14 is most concentrated in the superficial layers of sensory and motor cortices, but it is evenly distributed between superficial and deeper layers in association cortices (exemplified in Fig. 6I). Given that neurons in superficial layers are mainly involved in cortico-cortical communication, while deeper layer cells project to subcortical regions (Aboitiz, 1999), this pattern of localization may indicate that isocortical RGS14 functions regulate aspects of polymodal sensory processing, such as learning associations between polymodal stimuli and their capacity to drive behavior when experienced concurrently.

### Striatum

RGS14 labeling is broadly, but variably, seen throughout the striatum of WT mice, with “hotspots” of intense signal seen in nucleus accumbens core (NAcC, Fig. 7C & Fig. 8C). The position of these hotspots in Figure 1D, in consideration with the position of the thin band of intense signal presumed to be anterior commiussure in Figure 1E, suggests that these hotspots are likely at or near terminal processes of fibers traveling through the anterior commissure. This patchy protein distribution pattern is seen throughout the striatum, but generally becomes more diffuse as radial distance from the anterior commissure grows. In contrast to the high levels in NAcC, low levels of RGS14 immunoreactivity are seen in the surrounding nucleus accumbens shell (NAcS, Fig. 7C & Fig. 8C). Moderate levels are seen in the caudoputamen (CPu), with particular hotspots seen ventromedially throughout the entire rostrocaudal axis (Fig. 8C,F,I). Moderate levels are also seen in the fundus of the striatum (FS, Fig. 8F), a transitional region between CPu and pallidum, and low levels are seen in the olfactory tubercle (OT, Fig. 8C). Analysis of LR tissue revealed that RGS14-positive nuclei are found in a pattern that generally mirrors signal density in WT mice in all aforementioned subregions except for OT, in which nuclear RGS14 was not detected (Fig. 8B). Similar to SUBd, we suspect that RGS14 in OT is present in axons and terminals from projecting regions such as ENT and PERI.

**Fig. 7:**
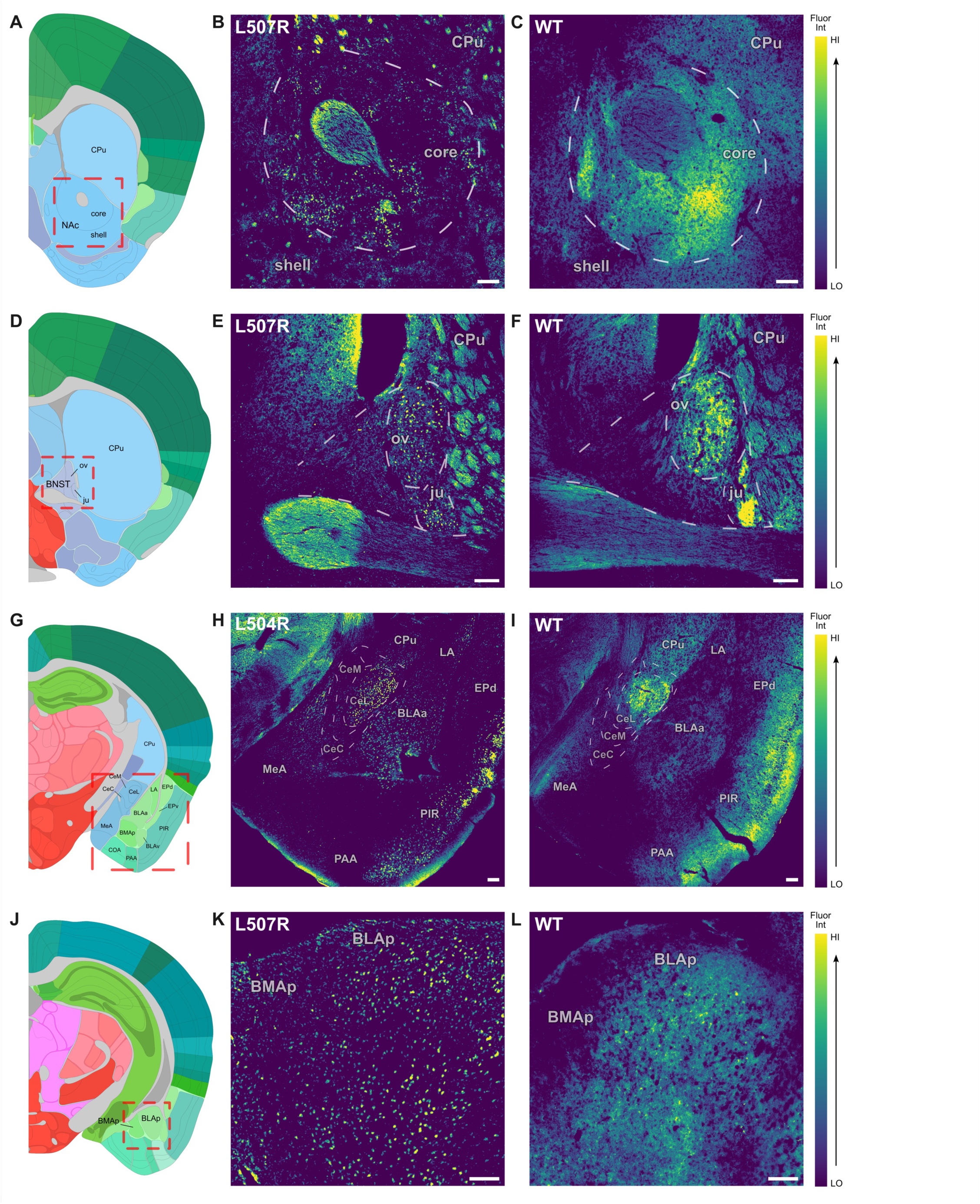
RGS14 is differentially expressed in distinct subregions of the amygdala, extended amygdala, and ventral striatum. Relative fluorescence intensity in RGS14-immunostained tissue sections from wild type (WT) and L507R-reporter mice. Anatomical annotations (left; **A,D,G,J**) from the Allen Mouse Brain Reference Atlas (atlas.brain-map.org) aligned with confocal images of RGS14 immunoreactivity in LR reporter (middle; **B,E,H,K**) and WT mice (right; **C,F,I,L**). Red dashed box in left panels indicates approximate anatomical location of tissue cross-sections in middle and right panels in each row. Scale bar = 100 µm. Abbreviations: **BLA(a,p,v)**: basolateral amygdala- (anterior, posterior, ventral), **BMAp**: basomedial amygdala-posterior, **BNST**: bed nucleus of the stria terminals, **Ce(C,L,M)**: central amygdala- (capsular, lateral, medial), **COA**: cortical amygdalar area, **CPu**: caudoputamen, **EP(d,v)**: endopiriform nucleus- (dorsal, ventral), **ju**: juxtacapsular nucleus of BNST, **LA**: lateral amygdala, **NAc**: nucleus accumbens, **MeA**: medial amygdala, **ov**: oval nucleus of the BNST, **PAA**: piriform-amygdalar area, **PIR**: piriform cortex.

**Fig. 8:**
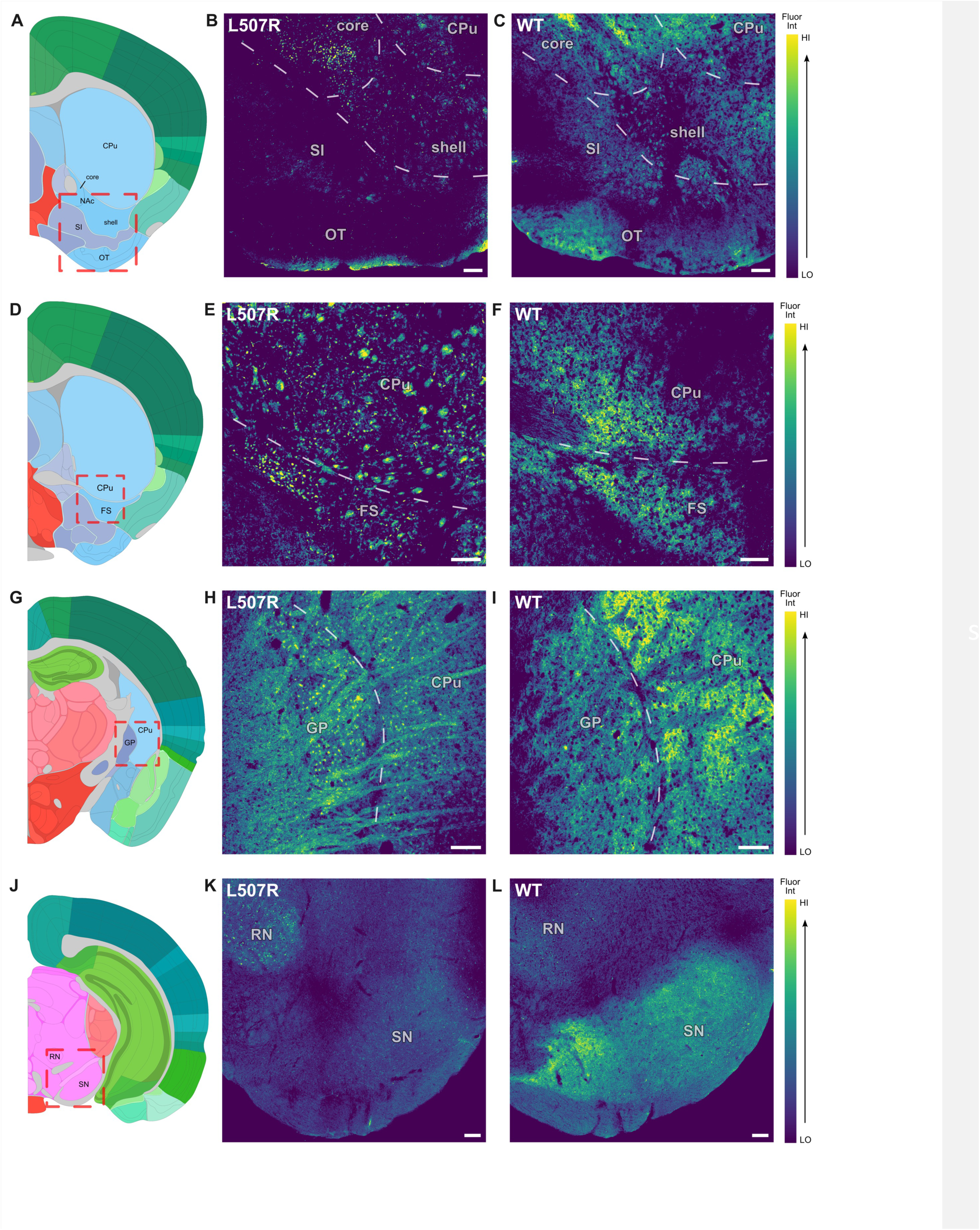
RGS14 is variably expressed across the basal ganglia. Relative fluorescence intensity in RGS14-immunostained tissue sections from wild type (WT) and L507R-reporter mice. Anatomical annotations (left; **A,D,G,J**) from the Allen Mouse Brain Reference Atlas (atlas.brain-map.org) aligned with confocal images of RGS14 immunoreactivity in LR reporter (middle; **B,E,H,K**) and WT mice (right; **C,F,I,L**). Red dashed box in left panels indicates approximate anatomical location of tissue cross-sections in middle and right panels in each row. Scale bar = 100 µm. Abbreviations: **CPu**: caudoputamen, **FS**: fundus of the striatum, **GP**: globus pallidus, **NAc**: nucleus accumbens, **OT**: olfactory tubercle, **RN**: red nucleus, **SI**: substantia innominata, **SN**: substantia nigra.

**Fig. 9:**
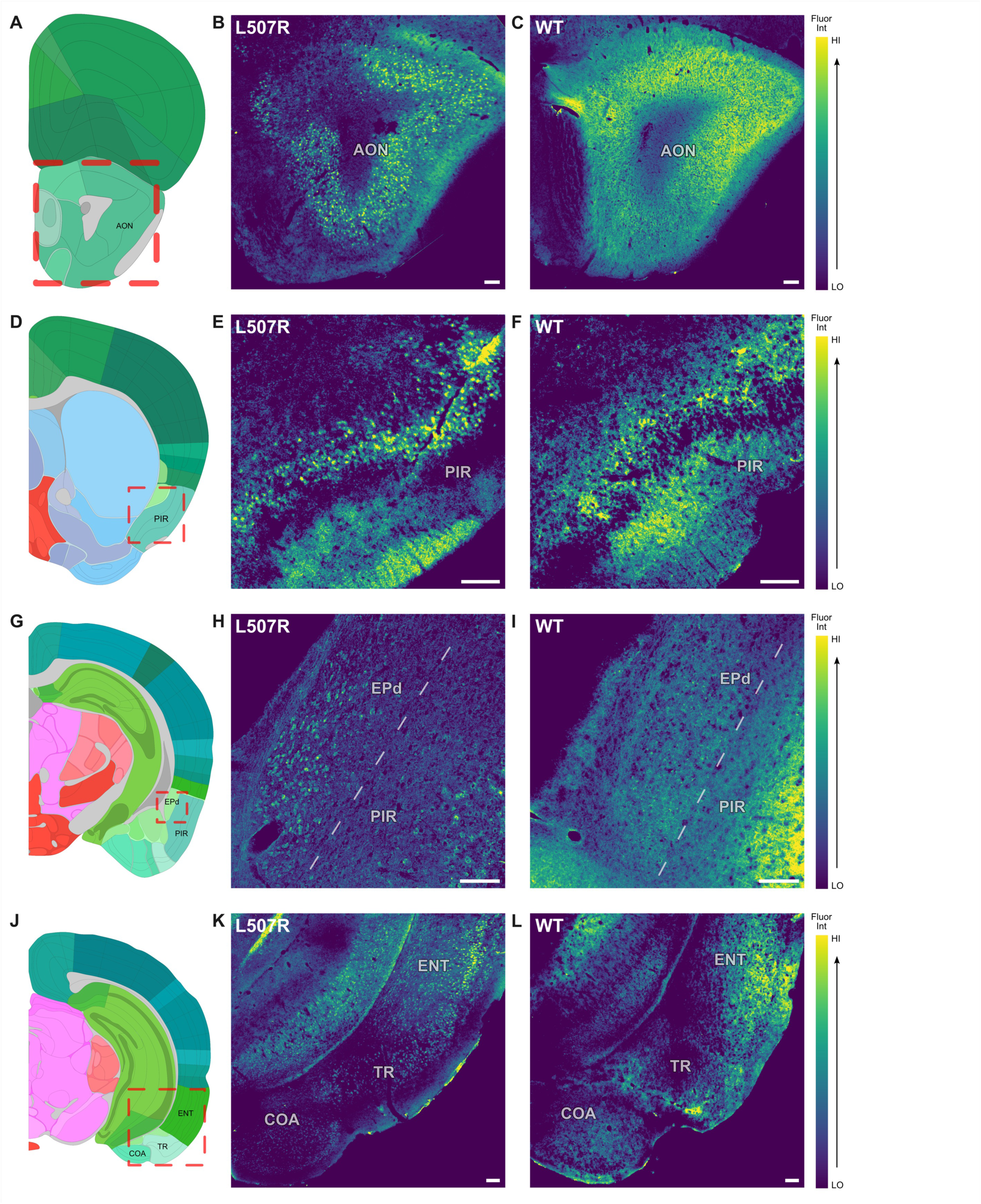
RGS14 is widely expressed at moderate levels across olfactory areas. Relative fluorescence intensity in RGS14-immunostained tissue sections from wild type (WT) and L507R-reporter mice. Anatomical annotations (left; **A,D,G,J**) from the Allen Mouse Brain Reference Atlas (atlas.brain-map.org) aligned with confocal images of RGS14 immunoreactivity in LR reporter (middle; **B,E,H,K**) and WT mice (right; **C,F,I,L**). Red dashed box in left panels indicates approximate anatomical location of tissue cross-sections in middle and right panels in each row. Scale bar = 100 µm. Abbreviations: **AON**: anterior olfactory nucleus, **COA**: cortical amygdalar area, **EPd**: endopiriform nucleus-dorsal, **ENT**: entorhinal cortex, **PIR**: piriform cortex, **TR**: postpiriform transition cortex.

### Pallidum and forebrain regions

RGS14 levels of immunostaining in the pallidum and neighboring regions are mostly light, except for two distinct hotspots in the bed nucleus of the stria terminalis (BNST) within the juxtacapsular nucleus (BNSTju) and oval nucleus (BNSTov, Fig. 7F). Analysis of the LR mice reveals densely-clustered RGS14-positive nuclei within both subregions in patterns that mirror signal density in WT mice (Fig. 7E). In neighboring regions, low signal is seen throughout the substantia innominata (SI, Fig. 8C), and light signal in the globus pallidus (GP, Fig. 8I). There appears to be moderate signal in the triangular nucleus of the septum (TRS), though there is likely some degree of autofluorescence contamination from the surrounding regions (Fig. 10I). LR tissue reveals that RGS14-positive nuclei are present in GP (Fig. 8H), but not detected in SI (Fig. 8B) or TRS (Fig. 10H). Again, we suggest that RGS14 is present in axonal and terminal processes in these regions.

**Fig. 10:**
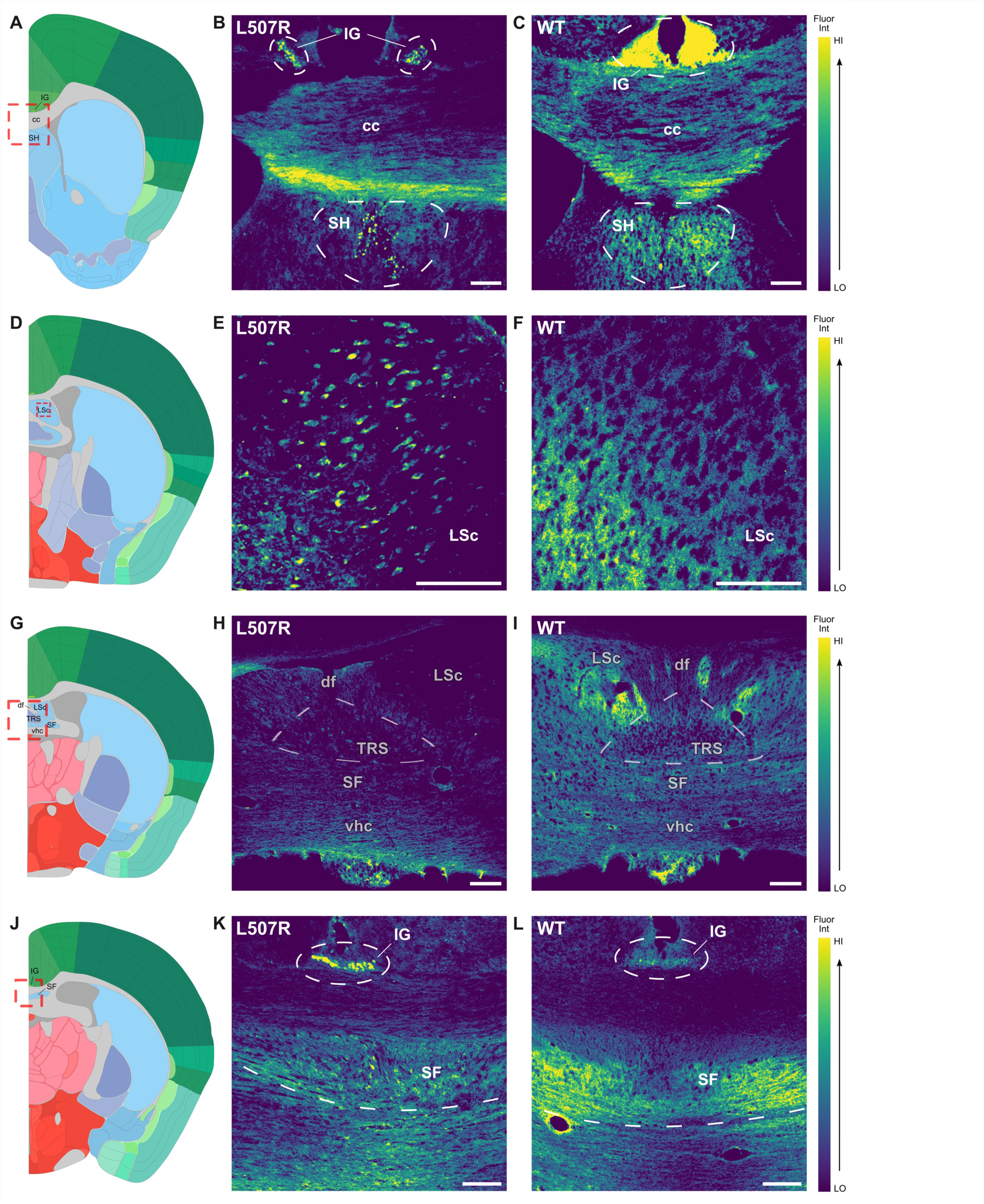
RGS14 is widely and intensely expressed in septal regions. Relative fluorescence intensity in RGS14-immunostained tissue sections from wild type (WT) and L507R-reporter mice. Anatomical annotations (left; **A,D,G,J**) from the Allen Mouse Brain Reference Atlas (atlas.brain-map.org) aligned with confocal images of RGS14 immunoreactivity in LR reporter (middle; **B,E,H,K**) and WT mice (right; **C,F,I,L**). Red dashed box in left panels indicates approximate anatomical location of tissue cross-sections in middle and right panels in each row. Scale bar = 100 µm. Abbreviations: **cc**: corpus callosum, **df**: dorsal fornix, **IG**: induseum griseum, **LSc**: lateral septal nucleus-caudal, **SF**: septofimbrial nucleus, **SH**: septohippocampal nucleus, **TRS**: triangular nucleus of the septum, **vhc**: ventral hippocampal commissure.

### Amygdala

The intensity of RGS14 protein labeling in the amygdala is regionally specific and mostly mild, with the exception of a hotspot in the lateral subdivision of the central amygdala (CeL). Signal in the surrounding medial and capsular subdivisions (CeM & CeC) is light, as is signal in the lateral amygdala (LA), anterodorsal portion of the medial amygdala (MeA), and anterior basolateral amygdala (BLAa, Fig. 7I) though it is somewhat more substantial in the posterior basolateral amygdala (BLAp, Fig. 7L). Analysis of LR tissue reveals that RGS14-positive nuclei are seen at a high density throughout the amygdala in a pattern that generally mirrors signal density in WT mice (Fig. 7H,K).

### Olfactory areas

In the olfactory areas, RGS14 immunoreactivity in WT mice is highest across ventrolateral allocortical regions, including the anterior olfactory nucleus (AON, Fig. 9C), piriform cortex (PIR, Fig. 7I & Fig. 9F), and postpiriform transition cortex (TR, Fig. 9L), where it is seen at moderate levels approximately evenly distributed been inner and outer layers. Staining in LR tissue shows RGS14-positive nuclei concentrated in the middle layer (AON: Fig. 9B, PIR: Fig. 7H & Fig. 9E), or the related inner layer for AON (Fig. 9B). RGS14 is present at low levels in select ventromedial allocortical regions, including the dorsal taenia tecta (TTd, Fig. 1B), nucleus of the lateral olfactory tract (NLOT, not shown), cortical amygdalar area (COA, Fig. 9L), and piriform-amygdalar area (PAA, Fig. 7I). LR tissue shows RGS14-positive nuclei in TTd (not shown) and COA (Fig. 9K), but not NLOT (not shown) or PAA (Fig. 7H), suggesting that RGS14 in NLOT and PAA is in axons and terminal processes. Lastly, light levels of RGS14 immunostaining are seen in the dorsal endopiriform nucleus (EPd, Fig. 9I). LR staining shows RGS14-positive nuclei in a pattern that generally mirrors signal density in WT mice (Fig. 9H).

### Midbrain

In WT mice, we found low-level RGS14 immunoreactivity in the substantia nigra (SN, Fig. 8L) that is most likely presynaptic given the absence of RGS14-positive nuclei in LR tissue in this region (Fig. 8K). This staining is found throughout the SN pars reticulata and possibly in the pars compacta as well, though it is difficult to say for sure without dopamine neurons labeling. We also found light staining in the red nucleus (RN, Fig. 8L), which is a component of descending dorsolateral motor pathways. The staining observed in LR tissue was unexpected, given that it is predominantly perinuclear instead of nuclear as expected (Fig. 8K). The reason for this surprising result is unknown, but a possibility is the existence of a regional splice variant in which the LR mutation instead interferes with nuclear import instead of nuclear export. It should be stated that some signal was noted at times in thalamic and hypothalamic regions in WT animals. We chose to exclude this from our results as we could not reliably distinguish it from background.

### Septal regions

The septum is a midline structure that consists of several limbic nuclei interwoven between fiber tracts of the fornix system. We show here that the septum as a whole is enriched in RGS14 immunoreactivity in both limbic nuclei and fiber tracts. Of the septal nuclei, the septofimbrial nucleus (SF) exhibits the most intense RGS14 immunoreactivity, especially in posterior portions (Fig. 10L). The caudodorsal subregion of the lateral septal nucleus (LSc) ranks closely behind SF in terms of RGS14 immunoreactivity, exhibiting high-level signal in this region (Fig, 10F,I), though signal is conspicuously absent in the rostroventral LS subregion. Tissue from LR mice shows RGS14-positive nuclei dispersed throughout SF (Fig. 10K) and LSc (Fig, 10E). The septohippocampal nucleus (SH) shows light levels of RGS14 staining in WT mice (Fig. 10C). LR tissue shows a band of RGS14-positive nuclei along the hemispheric midline within SH (Fig. 10B).

The fornix system tracts within the septum are strongly RGS14-immunoreactive as well (Fig. 1G & Fig. 2A). The murine dorsal fornix runs from the hippocampus along the septum, just ventral to the corpus collosum and medial to the SH (compare to Fig. 10B,C), following which it meets with the anterior commissure and disperses out towards its target nuclei. While tractology is complex and beyond the scope of this paper, it seems reasonable to suggest that these intra-septal tracts are a route through which RGS14 regulates communication between cortical and subcortical limbic regions.

## 4. DISCUSSION

RGS14 is a multifunctional signaling protein that serves as a natural suppressor of synaptic signaling and plasticity. Our results here present a comprehensive characterization of the distribution of RGS14 protein immunoreactivity throughout the adult mouse brain. The use of the RGS14 reporter mouse line (RGS14-LR) permitted us to positively identify RGS14 in regions that had previously escaped detection due to low or diffuse protein expression, and comparisons between LR and WT mice provided insight into key questions about the cellular origin and axonal expression of RGS14 in the brain. Overall, RGS14 immunoreactivity is found to varying degrees in cell bodies and neuropil in a variety of brain nuclei across the full extent of forebrain and midbrain regions in adult mice. The most intense levels are seen in the hippocampus proper, septum, NAcC subregion of the striatum, CeL subregion of the amygdala, and oval and juxtacapsular subregions of the BNST. It is also apparently trafficked along several major fiber tracts that involve cortico-limbic-basal ganglia connections that are central to emotional processing (Ingram, 2005; Willis & Haines, 2018). These results reaffirm and expand upon our previous knowledge of adult mouse brain RGS14 protein (Evans et al., 2014; Foster et al., 2021). Among the new discoveries are strong RGS14 immunoreactivity in discrete nuclei in the BNST and throughout nuclei and fiber tracts in the septum, and light immunoreactivity in isocortex that had previously escaped detection in adult mice.

We previously examined brain RGS14 expression in human and nonhuman primates and find that the murine expression profile largely aligns with what we know of it in primates with a few notable distinctions (Harbin et al., 2021; Montanez-Miranda et al., 2023; Squires et al., 2018). For one, it is differently distributed among the constituent subregions of the amygdala and striatum. In amygdala, both species show RGS14 hotspots in CeL, but murine RGS14 signal in BLA is significantly lower than in CeL and basomedial amygdala signal is very low if not undetectable, while staining in primate basomedial amygdala and BLA is just as strong as in CeL. Additionally, we show light signal in the LA in mice, while staining is reportedly absent in this region in macaques. In the striatum, murine NAc RGS14 labeling is higher than in CPu, while the reverse is true in primates. Another difference is seen in basal ganglia; RGS14 in murine GP appears to be localized pre- and post-synaptically, while evidence suggests that in the primate GP it is exclusively postsynaptic and is also seen at higher levels than in mouse.

Some notable features of its protein distribution profile hint to new roles for RGS14 in brain and behavior. RGS14 is known to suppress long-term potentiation and is required for long-term depression in hippocampal field CA2 (Evans et al., 2018b; Lee et al., 2010; Samadi et al., 2023). Our group has also recently shown that RGS14 protein expression can be markedly upregulated in the hippocampus as a protective mechanism in response to seizure-induced hyperactivity (Harbin et al., 2023). Considering that many RGS14-containing regions identified in this report are known to be conditionally plastic, it is possible that RGS14 functions as a tunable regulator of synaptic plasticity throughout many functionally related parts of the rodent brain. Here we will explore this possibility by highlighting several of the most noteworthy results and discussing the potential implications of regional RGS14 action on cognition, behavior, and human disease.

### RGS14 in the hippocampal formation and basolateral amygdala

RGS14 is abundant throughout the hippocampal formation in regions that mediate various components of learning and memory (Bett et al., 2013; Dimsdale-Zucker et al., 2018; Garcia & Buffalo, 2020; Hartopp et al., 2019; Jonas & Lisman, 2014; Park et al., 2022; Tzakis & Holahan, 2019). The implications of RGS14 in hippocampus proper have been previously discussed at length by our group and so will not be discussed here (Evans et al., 2014; Harbin et al., 2021; Montanez-Miranda et al., 2023). Regarding retrohippocampal regions, RGS14 is most abundant in ENT, which is critical for formation and recall of declarative memories due to its extensive relay between the cortex and the rest of the hippocampal formation (Canto et al., 2008; Garcia & Buffalo, 2020; Lee et al., 2021; Schultz et al., 2015). RGS14 is seen equally in medial ENT, which processes spatial memory and navigation, and lateral ENT, which processes associative learning of external stimuli (Lee et al., 2021). Previous studies have demonstrated that RGS14 suppresses medial ENT-dependent functions by showing that loss of RGS14 enhances spatial memory in the Morris water maze (Lee et al., 2010), as well as lateral ENT-dependent functions by showing that RGS14 loss enhances cued fear memory in female mice (Alexander et al., 2019). RGS14 is also abundant in SUB, which mediates episodic memory retrieval through connections with ENT and receives synapses from CA1 that undergo both long-term potentiation and long-term depression (Behr et al., 2009; Matsumoto et al., 2019; O’Mara et al., 2000; Roy et al., 2017)

RGS14 is found in limbic regions involved in processing both positively- and negatively-valenced emotions (Fadok et al., 2018; Pignatelli & Beyeler, 2019; Rizzi-Wise & Wang, 2021). It is found at high levels in LSc, which sits between field CA3 and the ventral tegmental area (VTA) in a tri-synaptic circuit that activates VTA dopamine neurons and drives the association of context with reward (Jiang et al., 2018). The BLA integrates incoming sensory and valuation inputs from temporal and prefrontal cortical regions, respectively, with previously-learned information about a given stimulus from the hippocampus to assign emotional salience to said stimulus (Fudge & Emiliano, 2003; Hintiryan et al., 2021). Valence-coded BLA neuronal populations drive appetitive or aversive behaviors through their outputs to other regions; for example, BLA-NAc projections are generally appetitive while BLA-CeA projections are generally aversive (Beyeler et al., 2016). Long-term potentiation of synapses onto BLA principal neurons has been shown to underlie emotion-driven associative learning and behavior (Goosens & Maren, 2002; C. Li & Rainnie, 2014; Rogan et al., 1997). As mentioned previously, RGS14 protein levels in murine BLA are low, but are much higher in primates (Squires et al., 2018). The reason for this species disparity is unknown, but could be related to the relative complexity of information conveyed by cortical inputs into the primate BLA compared to rodents (McDonald, 1998).

### RGS14 in the extended amygdala

The extended amygdala is a macrostructure consisting of interconnected regions with shared amygdala-like features from contiguous structures, namely CeA, BNST, NAcS, and SI. It is a key mediator of stress and threat responses, and is involved in certain types of emotional processing, particularly of negatively-valenced emotions such as fear and anxiety (Fox & Shackman, 2019). RGS14 is found throughout the extended amygdala, but distinct hotspots are observed in CeL, BNSTov, and BNSTju.

Information about fearful stimuli processed by the basolateral amygdala complex enters the central amygdala at CeL, is projected internally to CeC and CeM, and then exits the amygdala from CeM to the brainstem and hypothalamus to engage autonomic and neuroendocrine stress responses, and to the BNST (Pitkänen et al., 1997; Sah et al., 2003). CeL also sends long-range projections externally to the periaqueductal gray, which processes pain and defensive and aversive behaviors, and to the paraventricular nucleus of the thalamus, considered a critical node in the brain anxiety network (Kirouac, 2021; Penzo et al., 2014). Synaptic plasticity at local and long-range projection neurons in CeL is essential for fear learning, fear memory recall, and behavioral fear responses (Li et al., 2013; Penzo et al., 2014). Despite their many similarities in processing fearful stimuli, studies suggest that the central amygdala mediates the phasic emotional state that defines fear while the BNST mediates a sustained, lower-intensity fear response characteristic of anxiety (Ueda et al., 2021; Walker et al., 2009). Evidence exists for RGS14 involvement in both fear and anxiety behaviors based on fear conditioning and open field test studies (Alexander et al., 2019; Foster et al., 2021; Parker et al., 2012). The mutation used in these mouse studies was a global genetic deletion of RGS14 and as such the effects could not be definitively localized to specific brain regions, though one study found elevated c-Fos and pERK in CeA, CA1, and CA2 in RGS14-null mice following high-dose cocaine and novelty stress (Foster et al., 2021).

The BNSTov where RGS14 is expressed is tied to development of maladaptive behaviors in response to chronic and early life stress (Hu et al., 2020a; Hu et al., 2020b). Curiously, CeL and BNSTov produce the highest amounts of the stress hormone corticotropin-releasing factor (CRF) in the rodent brain outside of the hypothalamus (Hammack et al., 2021), and both regions are densely packed with RGS14. Neuropeptides such as CRF are capable of profound changes in the brain due to their prolonged duration of action and wide field of influence (Mantsch, 2022). It is possible that maintaining high basal levels of RGS14 in CeL and BNSTov exists as a preventative measure against over-recruitment of CRF systems. Future studies should examine RGS14 and CRF co-localization in these regions.

In contrast with BNSTov, the neighboring BNSTju is relatively under-studied but is also highly enriched with RGS14. The BNSTju has unique connections with the amygdala compared to other lateral BNST nuclei, including little to no innervation from central amygdala, dense glutamatergic innervation from BLA, and robust projections to CeM (H. Dong et al., 2000; H. W. Dong et al., 2001). BNSTju neurons experience long-term potentiation of intrinsic excitability (LTP-IE), which describes a form of non-synaptic plasticity characterized by a protracted decrease in firing threshold and increase in firing probability (Francesconi et al., 2009a; Francesconi et al., 2009b). It was determined that this occurs by inhibition of postsynaptic D-type K^+^ currents. RGS14 is packed extraordinarily densely into the miniscule BNSTju, though it is unknown what purpose this extreme expression is serving. Presumably RGS14 is suppressing plasticity in the region, but it is unclear if it is doing so canonically by disrupting synaptic plasticity onto BNSTju or by disrupting LTP-IE, or both (or perhaps it is doing another thing entirely). Protein tyrosine kinase activation has been shown to drive LTP-IE through D-type K+ channel internalization in CA3 pyramidal cells, and Ca^2+^/calmodulin-dependent kinase II (CaMKII) interacts with K+ channels to alter dendritic excitability in CA1 pyramidal cells (Hyun et al., 2013; Xu et al., 2005). RGS14 binds effectors from both H-Ras/ERK and Ca^++^/CaM/CaMKII signaling pathways, eluding to two mechanistic possibilities for interactions with this type of plasticity (Evans et al., 2018a; Shu et al., 2010; Willard et al., 2009). Future studies should explore RGS14 function in BNSTju, including the intriguing possible relationship between RGS14 and LTP-IE in BNSTju neurons.

### RGS14 in the striatum

The striatum is a large subcortical structure involved in processing goal-directed behavior and reward (De Groote & de Kerchove d’Exaerde, 2021). It serves as a nexus for the integration of signals from the cortex, limbic system, and brainstem into a unified signal that travels outward toward the thalamus through the basal ganglia (Young et al., 2024). The striatum is anatomically divided into ventral (NAcC and NAcS) and dorsal (CPu) regions, which process reward/aversion and conscious motor behaviors, respectively. The striatum and midbrain send reciprocal projections, forming hierarchical loops that shift the locus of control over behavior from NAcS → NAcC → CPu as a behavior transitions from volitional to habitual (Belin & Everitt, 2008; Haber et al., 2000). The densest striatal RGS14 signal is seen in the NAcC of the ventral striatum, a principal function of which is forming associations between stimuli and rewarding or punishing outcomes (Ambroggi et al., 2011; Cerri et al., 2014; Hamel et al., 2017; Ma et al., 2020; West & Carelli, 2016). RGS14 levels are more modest in NAcS, but are highest in the lateral portion which mediates reward seeking (Plasse et al., 2012; Smedley et al., 2019). Its dorsal striatal levels are also appreciable, most notably in ventral CPu. The ventral CPu receives various sensorimotor cortical inputs similar to the rest of the dorsal striatum, but it is uniquely connected to AgI, which is critically involved in salience processing (Gogolla, 2017; Hintiryan et al., 2016). As a whole, this protein distribution pattern points to a role of RGS14 in reward-related behavior. It has the capacity to impact multiple points of processing reward behavior in the striatum alone, including motivation for reward (lateral NAcS), which stimuli signal reward (NAcC), and execution of behavior to acquire reward (CPu).

While the reason for the opposite distribution of RGS14 in ventral and dorsal striatum in primate and mouse is unknown, we can speculate. Presuming that RGS14 is functioning to suppess plasticity, NAc RGS14 may act as a barrier to behavioral flexibility. This makes sense from an evolutionary perspective. In general, organisms predominantly engage in established behaviors because they promote continued survival, meaning that deviating from them can be risky. Risk, however, can lead to reward. Increased cognitive complexity lends itself to more complicated risk-reward calculations, which conceivably renders the capacity to readily adapt behavior more useful in highly complex organisms such as primates.

In this framework, as a barrier to plasticity, high NAc RGS14 levels could be adaptive in mice by favoring maintainence of established “safe” behaviors. Oppositely, less inhibitory tone on plasticity in NAc because of lower RGS14 levels could be adaptive in primates by favoring behavioral flexibility. This could also explain why RGS14 is more abundant in caudate and putamen in primate than in the homologous CPu in mice. High RGS14 levels in primate dorsal striatum could be suppressing habit formation, which also supports behavioral flexibility.

### Overview of RGS14 expression in mouse brain

The expression profile of RGS14 in the brain points to a possible role in governing aspects of several core cognitive functions: sensory perception, emotion, memory, motivation, and execution of actions. It is also in or just upstream of regions that process stress, which can have profound effects on cognitive functions (Marin et al., 2011). It seems that RGS14 may be particularly important to associative learning, which serves as a thread to tie many of these functions together. RGS14 is found in regions that form associations between concurrent stimuli (isocortex layer II), and between stimuli and context/content (ENT-hippocampus), social relationships (CA2), emotional states (BLA), responses (CPu), and outcomes (NAcC) (Beyeler et al., 2016; Cerri et al., 2014; J. Y. Lee et al., 2021; Samadi et al., 2023; Stalnaker et al., 2010; Wang & Cui, 2018). It may be telling that the regions in which RGS14 is most abundant are also especially dopaminoceptive (Hall et al., 1994). Dopamine is a strong driver of experience-based neuroplasticity, and RGS14 likely works to keep its influence in check (Speranza et al., 2021). While speculative, it is possible that RGS14 could even be induced in response to dopamine-driven hyperexcitation, similar to that which is seen during status epilepticus (Harbin et al., 2023).

Complex organisms are receiving and processing a multitude of stimuli every moment they are conscious. RGS14-mediated suppression of intracellular plasticity pathways could help filter the signal from the noise to improve the fidelity of associative learning. And because RGS14-expressing regions are interconnected, RGS14 may contribute to setting an overall tone on the experience-to-action processing pipeline that underlies emotional and motivated behaviors. Mutations that affect RGS14 function universally could have profound effects on attributes such as emotional resilience and emotional reactivity, compulsivity, impulsivity, and emotional memory. This is not just a hypothetical; the L507R mutation in the mice used in this study is a homolog of the naturally-occurring L505R genetic variant found in RGS14 of humans (Squires et al., 2020).

Addiction, post-traumatic stress disorder, and anxiety disorders (especially social anxiety) are just a few examples of the numerous psychopathologies that involve dysregulation of circuits with multiple RGS14-containing nodes. Among the brain regions that are changed in addiction are the striatum, amygdala, hippocampus, pallidum, and prefrontal cortex, all of which are RGS14-rich (Breiter et al., 1997; Childress et al., 2008; Gu et al., 2010; London et al., 1990; Sinha et al., 2005; Vollstädt-Klein et al., 2010). Additionally, abnormal function of the main CRF receptor in the brain predicts alcohol use disorder, and RGS14 is exceptionally densely packed into the highest extrahypothalamic CRF-producing regions in the brain (Blomeyer et al., 2008). Post-traumatic stress disorder is accompanied by decreases in hippocampal volume and increases in amygdala volume, and several types of anxiety disorders are linked to increased amygdala volume or connectivity (Bremner et al., 1995, 1997; Gurvits et al., 1996; Rauch et al., 1996; Redlich et al., 2015; Shin et al., 2004; Suor et al., 2020). Data regarding roles for BNST in humans is scarce, but it is likely that BNSTov RGS14 dysfunction could disrupt regulation of stress systems in ways that underlie and/or exacerbate pathological behaviors.

### Caveats and limitations

Immunofluorescent staining of brain tissue sections has some drawbacks mostly related to non-specific fluorescence, including issues with residual fluorescence near immunoreactivity hotspots, and autofluorescence generated by light scatter throughout tissue structural elements Also, fluorescence contamination from neuropil RGS14 reduces the resolution of immunofluorescence staining in WT mice, obscuring ultrastructural features such as cell bodies. This is a limitation that we partially address using the LR reporter mouse, in which RGS14 gets sequestered into cell nuclei and out of neuropil, dramatically increasing resolution specifically of RGS14-expressing cell bodies. Immunohistochemical approaches in general are vulnerable to non-specific edge stain artifacts, which are especially problematic for proteins localized to superficial cortical layers like RGS14. Conveniently, LR reporters address this problem as well by: 1) positively identifying RGS14-expressing nuclei in superficially-projecting cortical layers, and 2) serving as a secondary, pseudo-negative control for WT neuropil and cell body elements.

Another limitation of our approach stems from the semi-quantitive nature of relative fluorescence intensity as a measure of protein level, which is an approximation and limits the types of observations that can be made. This is further limited by the fact that atlas images could only be pseudo-standardized based on features like autofluorescence intensity in common tissue features, which is less reliable than, say, staining for an additional protein marker with established tissue protein levels to standardize to. An important caveat of our approach relates to the drawing of regional boundaries in microscopy images. Boundaries were manually drawn largely based on subjective matching of tissue structural features and fluorescence patterns to atlas images. While they are informed approximations of anatomical boundaries, they are inherently subjective and vulnerable to bias.

### Conclusions

In summary, our findings here demonstrate the broad expression profile of RGS14 in adult mouse brain and open new avenues for future studies to explore RGS14 actions in these interconnected brain circuits.

## ACKNOWLEDGMENTS

The authors would like to thank Dr. Daniel Lustberg for helpful discussions that inspired this project. This work was supported by the Emory University Integrated Cellular Imaging Core Facility (RRID:SCR_023534), NIH/HHS grant R01 NS037112 (awarded to JRH), and NIH/ORIP base grant of the Emory National Primate Research Center P51OD011132 (awarded to YS).

## ABBREVIATIONS

ACC: anterior cingulate cortex
AUD: auditory cortex
AgI: agranular insular cortex
AON: anterior olfactory nucleus
BLA(a,p): basolateral amygdala- (anterior, posterior)
BNST(ju,ov): bed nucleus of the stria terminals- (juxtacapsular, oval)
CA(1,2,3): hippocampal field *cornu ammonis* (1, 2,3)
CaM: calmodulin
CaMKII: calmodulin-dependent kinase II
CeA: central amygdala
Ce(C,L,M): central amygdala- (capsular, lateral, medial)
COA: cortical amygdalar area
CPu: caudoputamen
CRF: corticotropin-releasing factor
DG: dentate gyrus
dHIPP: dorsal hippocampus
ECT: ectorhinal cortex
ENT(l,m): entorhinal cortex- (lateral, medial)
EPd: endopiriform nucleus-dorsal
ERK: extracellular signal-regulated kinase
FC: fasciola cinerea
FS: fundus of the striatum
GP: globus pallidus
IG: induseum griseum
ju: juxtacapsular nucleus of the BNST
LA: lateral amygdala
LR: L507R mutant
LSc: lateral septal nucleus-caudal
LUT: look-up table
LTP-IE: long-term potentiation of intrinsic excitability
MeA: medial amygdala
MO: motor cortex
NAc(C,S): nucleus accumbens- (core, shell)
NGS: normal goat serum
NLOT: nucleus of the lateral olfactory tract
OT: olfactory tubercle
ov: oval nucleus of the BNST
PAA: piriform-amygdalar area
PBS: phosphate-buffered saline
PERI: perirhinal cortex
PFA: paraformaldehyde
PIR: piriform cortex
PL: prelimbic cortex
POST: postsubiculum
PTL: posterior parietal association cortex
RGS(14): regulator of G protein signaling (14)
RN: red nucleus
RSPv: retrosplenial cortex-ventral
SF: septofimbrial nucleus
SH: septohippocampal nucleus
SI: substantia innominata
SN(c,r): substantia nigra- (pars compacta, pars reticulata)
SS: somatosensory cortex
SUB(d,v): subiculum- (dorsal, ventral)
TeA: temporal association cortex
TR: postpiriform transition cortex
TRS: triangular nucleus of the septum
TTd: taenia tecta-dorsal
vHIPP: ventral hippocampus
VIS: visual cortex
VISC: visceral cortex
VTA: ventral tegmental area
WT: wild-type

## DATA AVAILABILITY

The raw data that support the findings of this study are available from the corresponding authors, S.N.B. and J.R.H, upon reasonable request.

## AUTHOR CONTRIBUTIONS

*Conceptualization*, S.N.B., N.H.H., and J.R.H.; *Methodology,* S.N.B., N.H.H., and J.R.H.; *Data collection*, S.N.B. and S.M.F., *Data organization*, S.N.B., W.Y., and C.B.; *Formal Analysis*, S.N.B., S.M.F., and N.H.H.; *Resources,* J.R.H.; *Literature Review*, S.N.B., W.Y., and G.V.D.; *Writing–Original Draft*, S.N.B. and J.R.H.; *Writing–Review & Editing*, S.N.B., Y.S., and J.R.H.; *Visualization*, S.N.B. and S.M.F.; *Supervision,* S.N.B., N.H.H., and J.R.H.; *Funding Acquisition*, J.R.H. All authors have read and agreed to the published version of the manuscript.

## REFERENCES

Aboitiz, F. (1999). Feature Article: Comparative Development of the Mammalian Isocortex and the Reptilian Dorsal Ventricular Ridge. Evolutionary Considerations. Cerebral Cortex, 9(8), 783–791. 10.1093/cercor/9.8.783

Agudelo, L. Z., Ferreira, D. M. S., Cervenka, I., Bryzgalova, G., Dadvar, S., Jannig, P. R., Pettersson-Klein, A. T., Lakshmikanth, T., Sustarsic, E. G., Porsmyr-Palmertz, M., Correia, J. C., Izadi, M., Martínez-Redondo, V., Ueland, P. M., Midttun, Ø., Gerhart-Hines, Z., Brodin, P., Pereira, T., Berggren, P.-O., & Ruas, J. L. (2018). Kynurenic Acid and Gpr35 Regulate Adipose Tissue Energy Homeostasis and Inflammation. Cell Metabolism, 27(2), 378–392.e5. 10.1016/j.cmet.2018.01.004

Alexander, G. M., Riddick, N. V., McCann, K. E., Lustberg, D., Moy, S. S., & Dudek, S. M. (2019). Modulation of CA2 neuronal activity increases behavioral responses to fear conditioning in female mice. Neurobiology of Learning and Memory, 163, 107044. 10.1016/j.nlm.2019.107044

Allen Institute for Brain Science (2011). Allen Reference Atlas – Mouse Brain. Available from atlas.brain-map.org.

Ambroggi, F., Ghazizadeh, A., Nicola, S. M., & Fields, H. L. (2011). Roles of Nucleus Accumbens Core and Shell in Incentive-Cue Responding and Behavioral Inhibition. The Journal of Neuroscience, 31(18), 6820–6830. 10.1523/JNEUROSCI.6491-10.2011

Behr, J., Wozny, C., Fidzinski, P., & Schmitz, D. (2009). Synaptic plasticity in the subiculum. Progress in Neurobiology, 89(4), 334–342. 10.1016/j.pneurobio.2009.09.002

Belin, D., & Everitt, B. J. (2008). Cocaine Seeking Habits Depend upon Dopamine-Dependent Serial Connectivity Linking the Ventral with the Dorsal Striatum. Neuron, 57(3), 432–441. 10.1016/j.neuron.2007.12.019

Bett, D., Stevenson, C. H., Shires, K. L., Smith, M. T., Martin, S. J., Dudchenko, P. A., & Wood, E. R. (2013). The Postsubiculum and Spatial Learning: The Role of Postsubicular Synaptic Activity and Synaptic Plasticity in Hippocampal Place Cell, Object, and Object-Location Memory. Journal of Neuroscience, 33(16), 6928–6943. 10.1523/JNEUROSCI.5476-12.2013

Beyeler, A., Namburi, P., Glober, G. F., Simonnet, C., Calhoon, G. G., Conyers, G. F., Luck, R., Wildes, C. P., & Tye, K. M. (2016). Divergent Routing of Positive and Negative Information from the Amygdala during Memory Retrieval. Neuron, 90(2), 348–361. 10.1016/j.neuron.2016.03.004

Blomeyer, D., Treutlein, J., Esser, G., Schmidt, M. H., Schumann, G., & Laucht, M. (2008). Interaction between CRHR1 gene and stressful life events predicts adolescent heavy alcohol use. Biological Psychiatry, 63(2), 146–151. 10.1016/j.biopsych.2007.04.026

Branch, M. R., & Hepler, J. R. (2017). Endogenous RGS14 is a cytoplasmic-nuclear shuttling protein that localizes to juxtanuclear membranes and chromatin-rich regions of the nucleus. PLoS ONE, 12(9), 1–27. 10.1371/journal.pone.0184497

Breiter, H. C., Gollub, R. L., Weisskoff, R. M., Kennedy, D. N., Makris, N., Berke, J. D., Goodman, J. M., Kantor, H. L., Gastfriend, D. R., Riorden, J. P., Mathew, R. T., Rosen, B. R., & Hyman, S. E. (1997). Acute effects of cocaine on human brain activity and emotion. Neuron, 19(3), 591–611. 10.1016/s0896-6273(00)80374-8

Bremner, J. D., Randall, P., Scott, T. M., Bronen, R. A., Seibyl, J. P., Southwick, S. M., Delaney, R. C., McCarthy, G., Charney, D. S., & Innis, R. B. (1995). MRI-Based Measurement of Hippocampal Volume in Patients With Combat-Related Posttraumatic Stress Disorder. The American Journal of Psychiatry, 152(7), 973. 10.1176/ajp.152.7.973

Bremner, J. D., Randall, P., Vermetten, E., Staib, L., Bronen, R. A., Mazure, C., Capelli, S., McCarthy, G., Innis, R. B., & Charney, D. S. (1997). Magnetic Resonance Imaging-Based Measurement of Hippocampal Volume in Posttraumatic Stress Disorder Related to Childhood Physical and Sexual Abuse—A Preliminary Report. Biological Psychiatry, 41(1), 23–32.

Canto, C. B., Wouterlood, F. G., & Witter, M. P. (2008). What Does the Anatomical Organization of the Entorhinal Cortex Tell Us? Neural Plasticity, 2008, e381243. 10.1155/2008/381243

Cerri, D. H., Saddoris, M. P., & Carelli, R. M. (2014). Nucleus Accumbens Core Neurons Encode Value-Independent Associations Necessary for Sensory Preconditioning. Behavioral Neuroscience, 128(5), 567–578. 10.1037/a0037797

Childress, A. R., Ehrman, R. N., Wang, Z., Li, Y., Sciortino, N., Hakun, J., Jens, W., Suh, J., Listerud, J., Marquez, K., Franklin, T., Langleben, D., Detre, J., & O’Brien, C. P. (2008). Prelude to Passion: Limbic Activation by “Unseen” Drug and Sexual Cues. PLoS ONE, 3(1), e1506. 10.1371/journal.pone.0001506

Cho, H., Kim, D.-U., & Kehrl, J. H. (2005). RGS14 Is a Centrosomal and Nuclear Cytoplasmic Shuttling Protein That Traffics to Promyelocytic Leukemia Nuclear Bodies Following Heat Shock *. Journal of Biological Chemistry, 280(1), 805–814. 10.1074/jbc.M408163200

De Groote, A., & de Kerchove d’Exaerde, A. (2021). Thalamo-Nucleus Accumbens Projections in Motivated Behaviors and Addiction. Frontiers in Systems Neuroscience, 15. 10.3389/fnsys.2021.711350

Dimsdale-Zucker, H. R., Ritchey, M., Ekstrom, A. D., Yonelinas, A. P., & Ranganath, C. (2018). CA1 and CA3 differentially support spontaneous retrieval of episodic contexts within human hippocampal subfields. Nature Communications, 9(1), 294. 10.1038/s41467-017-02752-1

Dong, H., Petrovich, G. D., & Swanson, L. W. (2000). Organization of projections from the juxtacapsular nucleus of the BST: A PHAL study in the rat. Brain Research, 859(1), 1–14. 10.1016/s0006-8993(99)02246-5

Dong, H. W., Petrovich, G. D., & Swanson, L. W. (2001). Topography of projections from amygdala to bed nuclei of the stria terminalis. Brain Research. Brain Research Reviews, 38(1–2), 192–246. 10.1016/s0165-0173(01)00079-0

Evans, P. R., Gerber, K. J., Dammer, E. B., Duong, D. M., Goswami, D., Lustberg, D. J., Zou, J., Yang, J. J., Dudek, S. M., Griffin, P. R., Seyfried, N. T., & Hepler, J. R. (2018a). Interactome Analysis Reveals Regulator of G Protein Signaling 14 (RGS14) is a Novel Calcium/Calmodulin (Ca 2+ /CaM) and CaM Kinase II (CaMKII) Binding Partner. Journal of Proteome Research, 17(4), 1700–1711. 10.1021/acs.jproteome.8b00027

Evans, P. R., Lee, S. E., Smith, Y., & Hepler, J. R. (2014). Postnatal developmental expression of regulator of G protein signaling 14 (RGS14) in the mouse brain. Journal of Comparative Neurology, 522(1), 186–203. 10.1002/cne.23395

Evans, P. R., Parra-Bueno, P., Smirnov, M. S., Lustberg, D. J., Dudek, S. M., Hepler, J. R., & Yasuda, R. (2018b). RGS14 restricts plasticity in hippocampal CA2 by limiting postsynaptic calcium signaling. eNeuro, 5(3), 1–13. 10.1523/ENEURO.0353-17.2018

Fadok, J. P., Markovic, M., Tovote, P., & Lüthi, A. (2018). New perspectives on central amygdala function. Current Opinion in Neurobiology, 49, 141–147. 10.1016/j.conb.2018.02.009

Foster, S. L., Lustberg, D. J., Harbin, N. H., Bramlett, S. N., Hepler, J. R., & Weinshenker, D. (2021). RGS14 modulates locomotor behavior and ERK signaling induced by environmental novelty and cocaine within discrete limbic structures. Psychopharmacology, 238(10), 2755–2773. 10.1007/s00213-021-05892-x

Fox, A. S., & Shackman, A. J. (2019). The central extended amygdala in fear and anxiety: Closing the gap between mechanistic and neuroimaging research. Neuroscience Letters, 693, 58–67. 10.1016/j.neulet.2017.11.056

Francesconi, W., Berton, F., Koob, G. F., & Sanna, P. P. (2009a). Intrinsic neuronal plasticity in the juxtacapsular nucleus of the bed nuclei of the stria terminalis (jcBNST). Progress in Neuro-Psychopharmacology and Biological Psychiatry, 33(8), 1347–1355. 10.1016/j.pnpbp.2009.08.003

Francesconi, W., Berton, F., Repunte-Canonigo, V., Hagihara, K., Thurbon, D., Lekic, D., Specio, S. E., Greenwell, T. N., Chen, S. A., Rice, K. C., Richardson, H. N., O’Dell, L. E., Zorrilla, E. P., Morales, M., Koob, G. F., & Sanna, P. P. (2009b). Protracted withdrawal from alcohol and drugs of abuse impairs long-term potentiation of intrinsic excitability in the juxtacapsular bed nucleus of the stria terminalis. The Journal of Neuroscience: The Official Journal of the Society for Neuroscience, 29(17), 5389–5401. 10.1523/JNEUROSCI.5129-08.2009

Friedman, P. A., Sneddon, W. B., Mamonova, T., Montanez-Miranda, C., Ramineni, S., Harbin, N. H., Squires, K. E., Gefter, J. V., Magyar, C. E., Emlet, D. R., & Hepler, J. R. (2022). RGS14 regulates PTH- and FGF23-sensitive NPT2A-mediated renal phosphate uptake via binding to the NHERF1 scaffolding protein. The Journal of Biological Chemistry, 298(5), 101836. 10.1016/j.jbc.2022.101836

Fudge, J. L., & Emiliano, A. B. (2003). The Extended Amygdala and the Dopamine System: Another Piece of the Dopamine Puzzle. The Journal of Neuropsychiatry and Clinical Neurosciences, 15(3), 306–316.

Garcia, A. D., & Buffalo, E. A. (2020). Anatomy and Function of the Primate Entorhinal Cortex. Annual Review of Vision Science, 6(Volume 6, 2020), 411–432. 10.1146/annurev-vision-030320-041115

Gogolla, N. (2017). The insular cortex. Current Biology, 27(12), R580–R586. 10.1016/j.cub.2017.05.010

Goosens, K. A., & Maren, S. (2002). Long-term potentiation as a substrate for memory: Evidence from studies of amygdaloid plasticity and Pavlovian fear conditioning. Hippocampus, 12(5), 592–599. 10.1002/hipo.10099

Gu, H., Salmeron, B. J., Ross, T. J., Geng, X., Zhan, W., Stein, E. A., & Yang, Y. (2010). Mesocorticolimbic Circuits are Impaired in Chronic Cocaine Users as Demonstrated by Resting State Functional Connectivity. NeuroImage, 53(2), 593–601. 10.1016/j.neuroimage.2010.06.066

Gurvits, T. V., Shenton, M. E., Hokama, H., Ohta, H., Lasko, N. B., Gilbertson, M. W., Orr, S. P., Kikinis, R., Jolesz, F. A., McCarley, R. W., & Pitman, R. K. (1996). Magnetic resonance imaging study of hippocampal volume in chronic, combat-related posttraumatic stress disorder. Biological Psychiatry, 40(11), 1091–1099. 10.1016/S0006-3223(96)00229-6

Haber, S. N., Fudge, J. L., & McFarland, N. R. (2000). Striatonigrostriatal Pathways in Primates Form an Ascending Spiral from the Shell to the Dorsolateral Striatum. The Journal of Neuroscience, 20(6), 2369–2382. 10.1523/JNEUROSCI.20-06-02369.2000

Hall, H., Sedvall, G., Magnusson, O., Kopp, J., Halldin, C., & Farde, L. (1994). Distribution of D1- and D2-Dopamine Receptors, and Dopamine and Its Metabolites in the Human Brain. Neuropsychopharmacology, 11(4), 245–256. 10.1038/sj.npp.1380111

Hamel, L., Thangarasa, T., Samadi, O., & Ito, R. (2017). Caudal Nucleus Accumbens Core Is Critical in the Regulation of Cue-Elicited Approach-Avoidance Decisions. eNeuro, 4(1). 10.1523/ENEURO.0330-16.2017

Hammack, S. E., Braas, K. M., & May, V. (2021). Chemoarchitecture of the BNST: Neurophenotypic diversity and function. Handbook of Clinical Neurology, 179, 385–402. 10.1016/B978-0-12-819975-6.00025-X

Harbin, N. H., Bramlett, S. N., Montanez-Miranda, C., Terzioglu, G., & Hepler, J. R. (2021). RGS14 Regulation of Post-Synaptic Signaling and Spine Plasticity in Brain. International Journal of Molecular Sciences, 22(13). 10.3390/ijms22136823

Harbin, N. H., Lustberg, D. J., Hurst, C., Pare, J.-F., Crotty, K. M., Waters, A. L., Yeligar, S. M., Smith, Y., Seyfried, N. T., Weinshenker, D., & Hepler, J. R. (2023). RGS14 is neuroprotective against seizure-induced mitochondrial oxidative stress and pathology in hippocampus. bioRxiv: The Preprint Server for Biology, 2023.02.01.526349. 10.1101/2023.02.01.526349

Hartopp, N., Wright, P., Ray, N. J., Evans, T. E., Metzler-Baddeley, C., Aggleton, J. P., & O’Sullivan, M. J. (2019). A Key Role for Subiculum-Fornix Connectivity in Recollection in Older Age. Frontiers in Systems Neuroscience, 12. 10.3389/fnsys.2018.00070

Hintiryan, H., Bowman, I., Johnson, D. L., Korobkova, L., Zhu, M., Khanjani, N., Gou, L., Gao, L., Yamashita, S., Bienkowski, M. S., Garcia, L., Foster, N. N., Benavidez, N. L., Song, M. Y., Lo, D., Cotter, K. R., Becerra, M., Aquino, S., Cao, C.,… Dong, H.-W. (2021). Connectivity characterization of the mouse basolateral amygdalar complex. Nature Communications, 12(1), 2859. 10.1038/s41467-021-22915-5

Hintiryan, H., Foster, N. N., Bowman, I., Bay, M., Song, M. Y., Gou, L., Yamashita, S., Bienkowski, M. S., Zingg, B., Zhu, M., Yang, X. W., Shih, J. C., Toga, A. W., & Dong, H.-W. (2016). The mouse cortico-striatal projectome. Nature Neuroscience, 19(8), 1100– 1114. 10.1038/nn.4332

Hollinger, S., Taylor, J. B., Goldman, E. H., & Hepler, J. R. (2001). RGS14 is a bifunctional regulator of Gαi/o activity that exists in multiple populations in brain. Journal of Neurochemistry, 79(5), 941–949. 10.1046/j.1471-4159.2001.00629.x

Hu, P., Liu, J., Maita, I., Kwok, C., Gu, E., Gergues, M. M., Kelada, F., Phan, M., Zhou, J.-N., Swaab, D. F., Pang, Z. P., Lucassen, P. J., Roepke, T. A., & Samuels, B. A. (2020a). Chronic Stress Induces Maladaptive Behaviors by Activating Corticotropin-Releasing Hormone Signaling in the Mouse Oval Bed Nucleus of the Stria Terminalis. Journal of Neuroscience, 40(12), 2519–2537. 10.1523/JNEUROSCI.2410-19.2020

Hu, P., Maita, I., Phan, M. L., Gu, E., Kwok, C., Dieterich, A., Gergues, M. M., Yohn, C. N., Wang, Y., Zhou, J.-N., Qi, X.-R., Swaab, D. F., Pang, Z. P., Lucassen, P. J., Roepke, T. A., & Samuels, B. A. (2020b). Early-life stress alters affective behaviors in adult mice through persistent activation of CRH-BDNF signaling in the oval bed nucleus of the stria terminalis. Translational Psychiatry, 10(1), 1–17. 10.1038/s41398-020-01070-3

Hyun, J. H., Eom, K., Lee, K.-H., Ho, W.-K., & Lee, S.-H. (2013). Activity-dependent downregulation of D-type K+ channel subunit Kv1.2 in rat hippocampal CA3 pyramidal neurons. The Journal of Physiology, 591(22), 5525–5540. 10.1113/jphysiol.2013.259002

Ingram, C. D. (2005). Chapter 4.11—Pathways transmitter interactions mediating an integrated stress response. In T. Steckler, N. H. Kalin, & J. M. H. M. Reul (Eds.), Techniques in the Behavioral and Neural Sciences (Vol. 15, pp. 609–639). Elsevier. 10.1016/S0921-0709(05)80033-1

Jiang, J.-X., Liu, H., Huang, Z.-Z., Cui, Y., Zhang, X.-Q., Zhang, X.-L., Cui, Y., & Xin, W.-J. (2018). The role of CA3-LS-VTA loop in the formation of conditioned place preference induced by context-associated reward memory for morphine. Addiction Biology, 23(1), 41–54. 10.1111/adb.12468

Jonas, P., & Lisman, J. (2014). Structure, function, and plasticity of hippocampal dentate gyrus microcircuits. Frontiers in Neural Circuits, 8. 10.3389/fncir.2014.00107

Kirouac, G. J. (2021). The Paraventricular Nucleus of the Thalamus as an Integrating and Relay Node in the Brain Anxiety Network. Frontiers in Behavioral Neuroscience, 15. 10.3389/fnbeh.2021.627633

Lee, J. Y., Jun, H., Soma, S., Nakazono, T., Shiraiwa, K., Dasgupta, A., Nakagawa, T., Xie, J. L., Chavez, J., Romo, R., Yungblut, S., Hagihara, M., Murata, K., & Igarashi, K. M. (2021). Dopamine facilitates associative memory encoding in the entorhinal cortex. Nature, 598(7880), 321–326. 10.1038/s41586-021-03948-8

Lee, S. E., Simons, S. B., Heldt, S. A., Zhao, M., Schroeder, J. P., Vellano, C. P., Cowan, D. P., Ramineni, S., Yates, C. K., Feng, Y., Smith, Y., Sweatt, J. D., Weinshenker, D., Ressler, K. J., Dudek, S. M., & Hepler, J. R. (2010). RGS14 is a natural suppressor of both synaptic plasticity in CA2 neurons and hippocampal-based learning and memory. Proceedings of the National Academy of Sciences, 107(39), 16994–16998. 10.1073/pnas.1005362107

Li, C., & Rainnie, D. G. (2014). Bidirectional regulation of synaptic plasticity in the basolateral amygdala induced by the D1-like family of dopamine receptors and group II metabotropic glutamate receptors. The Journal of Physiology, 592(Pt 19), 4329–4351. 10.1113/jphysiol.2014.277715

Li, H., Penzo, M. A., Taniguchi, H., Kopec, C. D., Huang, Z. J., & Li, B. (2013). Experience-dependent modification of a central amygdala fear circuit. Nature Neuroscience, 16(3), 332–339. 10.1038/nn.3322

Li, Y., Tang, X., Li, X., Dai, H., Miao, R., Cai, J., Huang, Z., Chen, A. F., Xing, X., Lu, Y., & Yuan, H. (2016). Regulator of G protein signalling 14 attenuates cardiac remodelling through the MEK–ERK1/2 signalling pathway. Basic Research in Cardiology, 111, 47. 10.1007/s00395-016-0566-1

London, E. D., Cascella, N. G., Wong, D. F., Phillips, R. L., Dannals, R. F., Links, J. M., Herning, R., Grayson, R., Jaffe, J. H., & Wagner, H. N. (1990). Cocaine-induced reduction of glucose utilization in human brain. A study using positron emission tomography and [fluorine 18]-fluorodeoxyglucose. Archives of General Psychiatry, 47(6), 567–574. 10.1001/archpsyc.1990.01810180067010

Ma, L., Chen, W., Yu, D., & Han, Y. (2020). Brain-Wide Mapping of Afferent Inputs to Accumbens Nucleus Core Subdomains and Accumbens Nucleus Subnuclei. Frontiers in Systems Neuroscience, 14. 10.3389/fnsys.2020.00015

Mantsch, J. R. (2022). Corticotropin releasing factor and drug seeking in substance use disorders: Preclinical evidence and translational limitations. Addiction Neuroscience, 4, 100038. 10.1016/j.addicn.2022.100038

Marin, M.-F., Lord, C., Andrews, J., Juster, R.-P., Sindi, S., Arsenault-Lapierre, G., Fiocco, A. J., & Lupien, S. J. (2011). Chronic stress, cognitive functioning and mental health. Neurobiology of Learning and Memory, 96(4), 583–595. 10.1016/j.nlm.2011.02.016

Matsumoto, N., Kitanishi, T., & Mizuseki, K. (2019). The subiculum: Unique hippocampal hub and more. Neuroscience Research, 143, 1–12. 10.1016/j.neures.2018.08.002

McDonald, A. J. (1998). Cortical pathways to the mammalian amygdala. Progress in Neurobiology, 55(3), 257–332. 10.1016/S0301-0082(98)00003-3

Montanez-Miranda, C., Bramlett, S. N., & Hepler, J. R. (2023). RGS14 expression in CA2 hippocampus, amygdala, and basal ganglia: Implications for human brain physiology and disease. Hippocampus, 33(3), 166–181. 10.1002/hipo.23492

O’Mara, S. M., Commins, S., & Anderson, M. (2000). Synaptic plasticity in the hippocampal area CA1-subiculum projection: Implications for theories of memory. Hippocampus, 10(4), 447–456. 10.1002/1098-1063(2000)10:4<447::AID-HIPO11>3.0.CO;2-2

Park, S.-B., Lim, H.-Y., Lee, E.-Y., Yoo, S.-W., Jung, H.-S., Lee, E., Sun, W., & Lee, I. (2022). The fasciola cinereum subregion of the hippocampus is important for the acquisition of visual contextual memory. Progress in Neurobiology, 210, 102217. 10.1016/j.pneurobio.2022.102217

Parker, C. C., Sokoloff, G., Cheng, R., & Palmer, A. A. (2012). Genome-Wide Association for Fear Conditioning in an Advanced Intercross Mouse Line. Behavior Genetics, 42(3), 437–448. 10.1007/s10519-011-9524-8

Penzo, M. A., Robert, V., & Li, B. (2014). Fear Conditioning Potentiates Synaptic Transmission onto Long-Range Projection Neurons in the Lateral Subdivision of Central Amygdala. Journal of Neuroscience, 34(7), 2432–2437. 10.1523/JNEUROSCI.4166-13.2014

Pignatelli, M., & Beyeler, A. (2019). Valence coding in amygdala circuits. Current Opinion in Behavioral Sciences, 26, 97–106. 10.1016/j.cobeha.2018.10.010

Pitkänen, A., Savander, V., & LeDoux, J. E. (1997). Organization of intra-amygdaloid circuitries in the rat: An emerging framework for understanding functions of the amygdala. Trends in Neurosciences, 20(11), 517–523. 10.1016/S0166-2236(97)01125-9

Plasse, G. van der, Schrama, R., Seters, S. P. van, Vanderschuren, L. J. M. J., & Westenberg, H. G. M. (2012). Deep Brain Stimulation Reveals a Dissociation of Consummatory and Motivated Behaviour in the Medial and Lateral Nucleus Accumbens Shell of the Rat. PLOS ONE, 7(3), e33455. 10.1371/journal.pone.0033455

Rauch, S. L., van der Kolk, B. A., Fisler, R. E., Alpert, N. M., Orr, S. P., Savage, C. R., Fischman, A. J., Jenike, M. A., & Pitman, R. K. (1996). A symptom provocation study of posttraumatic stress disorder using positron emission tomography and script-driven imagery. Archives of General Psychiatry, 53(5), 380–387. 10.1001/archpsyc.1996.01830050014003

Redlich, R., Grotegerd, D., Opel, N., Kaufmann, C., Zwitserlood, P., Kugel, H., Heindel, W., Donges, U.-S., Suslow, T., Arolt, V., & Dannlowski, U. (2015). Are you gonna leave me? Separation anxiety is associated with increased amygdala responsiveness and volume. Social Cognitive and Affective Neuroscience, 10(2), 278–284. 10.1093/scan/nsu055

Rizzi-Wise, C. A., & Wang, D. V. (2021). Putting Together Pieces of the Lateral Septum: Multifaceted Functions and Its Neural Pathways. eNeuro, 8(6), ENEURO.0315-21.2021. 10.1523/ENEURO.0315-21.2021

Rogan, M. T., Stäubli, U. V., & LeDoux, J. E. (1997). Fear conditioning induces associative long-term potentiation in the amygdala. Nature, 390(6660), 604–607. 10.1038/37601

Roy, D. S., Kitamura, T., Okuyama, T., Ogawa, S. K., Sun, C., Obata, Y., Yoshiki, A., & Tonegawa, S. (2017). Distinct Neural Circuits for the Formation and Retrieval of Episodic Memories. Cell, 170(5), 1000–1012.e19. 10.1016/j.cell.2017.07.013

Sah, P., Faber, E. S. L., Lopez De Armentia, M., & Power, J. (2003). The Amygdaloid Complex: Anatomy and Physiology. Physiological Reviews, 83(3), 803–834. 10.1152/physrev.00002.2003

Samadi, M., Hales, C. A., Lustberg, D. J., Farris, S., Ross, M. R., Zhao, M., Hepler, J. R., Harbin, N. H., Robinson, E. S. J., Banks, P. J., Bashir, Z. I., & Dudek, S. M. (2023). Mechanisms of mGluR-dependent plasticity in hippocampal area CA2. Hippocampus, 33(6), 730–744. 10.1002/hipo.23529

Schultz, H., Sommer, T., & Peters, J. (2015). The Role of the Human Entorhinal Cortex in a Representational Account of Memory. Frontiers in Human Neuroscience, 9. 10.3389/fnhum.2015.00628

Shin, L. M., Orr, S. P., Carson, M. A., Rauch, S. L., Macklin, M. L., Lasko, N. B., Peters, P. M., Metzger, L. J., Dougherty, D. D., Cannistraro, P. A., Alpert, N. M., Fischman, A. J., & Pitman, R. K. (2004). Regional cerebral blood flow in the amygdala and medial prefrontal cortex during traumatic imagery in male and female Vietnam veterans with PTSD. Archives of General Psychiatry, 61(2), 168–176. 10.1001/archpsyc.61.2.168

Shu, F., Ramineni, S., & Hepler, J. R. (2010). RGS14 is a multifunctional scaffold that integrates G protein and Ras/Raf MAPkinase signalling pathways. Cellular Signalling, 22(3), 366–376. 10.1016/j.cellsig.2009.10.005

Sinha, R., Lacadie, C., Skudlarski, P., Fulbright, R. K., Rounsaville, B. J., Kosten, T. R., & Wexler, B. E. (2005). Neural activity associated with stress-induced cocaine craving: A functional magnetic resonance imaging study. Psychopharmacology, 183(2), 171–180. 10.1007/s00213-005-0147-8

Sjögren, B. (2011). 10 - Regulator of G Protein Signaling Proteins as Drug Targets: Current State and Future Possibilities. In R. R. Neubig (Ed.), Advances in Pharmacology (Vol. 62, pp. 315–347). Academic Press. 10.1016/B978-0-12-385952-5.00002-6

Smedley, E. B., DiLeo, A., & Smith, K. S. (2019). Circuit directionality for motivation: Lateral accumbens-pallidum, but not pallidum-accumbens, connections regulate motivational attraction to reward cues. Neurobiology of Learning and Memory, 162, 23–35. 10.1016/j.nlm.2019.05.001

Speranza, L., di Porzio, U., Viggiano, D., de Donato, A., & Volpicelli, F. (2021). Dopamine: The Neuromodulator of Long-Term Synaptic Plasticity, Reward and Movement Control. Cells, 10(4), 735. 10.3390/cells10040735

Squires, K. E., Gerber, K. J., Pare, J.-F., Branch, M. R., Smith, Y., & Hepler, J. R. (2018). Regulator of G protein signaling 14 (RGS14) is expressed pre- and postsynaptically in neurons of hippocampus, basal ganglia, and amygdala of monkey and human brain. Brain Structure and Function, 223(1), 233–253. 10.1007/s00429-017-1487-y

Squires, K. E., Gerber, K. J., Tillman, M. C., Lustberg, D. J., Montañez-Miranda, C., Zhao, M., Ramineni, S., Scharer, C. D., Saha, R. N., Shu, F.-J., Schroeder, J. P., Ortlund, E. A., Weinshenker, D., Dudek, S. M., & Hepler, J. R. (2020). Human genetic variants disrupt RGS14 nuclear shuttling and regulation of LTP in hippocampal neurons. The Journal of Biological Chemistry, 296, 100024. 10.1074/jbc.RA120.016009

Stalnaker, T. A., Calhoon, G. G., Ogawa, M., Roesch, M. R., & Schoenbaum, G. (2010). Neural Correlates of Stimulus–Response and Response–Outcome Associations in Dorsolateral Versus Dorsomedial Striatum. Frontiers in Integrative Neuroscience, 4, 12. 10.3389/fnint.2010.00012

Stewart, A., & Fisher, R. A. (2015). Chapter One - Introduction: G Protein-coupled Receptors and RGS Proteins. In R. A. Fisher (Ed.), Progress in Molecular Biology and Translational Science (Vol. 133, pp. 1–11). Academic Press. 10.1016/bs.pmbts.2015.03.002

Suor, J. H., Jimmy, J., Monk, C. S., Phan, K. L., & Burkhouse, K. L. (2020). Parsing Differences in Amygdala Volume Among Individuals with and without Social and Generalized Anxiety Disorders Across the Lifespan. Journal of Psychiatric Research, 128, 83–89. 10.1016/j.jpsychires.2020.05.027

Tzakis, N., & Holahan, M. R. (2019). Social Memory and the Role of the Hippocampal CA2 Region. Frontiers in Behavioral Neuroscience, 13. 10.3389/fnbeh.2019.00233

Ueda, S., Hosokawa, M., Arikawa, K., Takahashi, K., Fujiwara, M., Kakita, M., Fukada, T., Koyama, H., Horigane, S., Itoi, K., Kakeyama, M., Matsunaga, H., Takeyama, H., Bito, H., & Takemoto-Kimura, S. (2021). Distinctive Regulation of Emotional Behaviors and Fear-Related Gene Expression Responses in Two Extended Amygdala Subnuclei With Similar Molecular Profiles. Frontiers in Molecular Neuroscience, 14, 741895. 10.3389/fnmol.2021.741895

Vellano, C. P., Brown, N. E., Blumer, J. B., & Hepler, J. R. (2013). Assembly and function of the regulator of G protein signaling 14 (RGS14)·H-Ras signaling complex in live cells are regulated by Gαi1 and Gαi-linked G protein-coupled receptors. The Journal of Biological Chemistry, 288(5), 3620–3631. 10.1074/jbc.M112.440057

Vellano, C. P., Maher, E. M., Hepler, J. R., & Blumer, J. B. (2011). G protein-coupled receptors and resistance to inhibitors of cholinesterase-8A (Ric-8A) both regulate the regulator of g protein signaling 14 RGS14·Gαi1 complex in live cells. The Journal of Biological Chemistry, 286(44), 38659–38669. 10.1074/jbc.M111.274928

Vollstädt-Klein, S., Wichert, S., Rabinstein, J., Bühler, M., Klein, O., Ende, G., Hermann, D., & Mann, K. (2010). Initial, habitual and compulsive alcohol use is characterized by a shift of cue processing from ventral to dorsal striatum. Addiction, 105(10), 1741–1749. 10.1111/j.1360-0443.2010.03022.x

Walker, D. L., Miles, L. A., & Davis, M. (2009). Selective Participation of the Bed Nucleus of the Stria Terminalis and CRF in Sustained Anxiety-Like versus Phasic Fear-Like Responses. Progress in Neuro-Psychopharmacology & Biological Psychiatry, 33(8), 1291–1308. 10.1016/j.pnpbp.2009.06.022

Wang, J.-H., & Cui, S. (2018). Associative memory cells and their working principle in the brain. F1000Research, 7, 108. 10.12688/f1000research.13665.1

West, E. A., & Carelli, R. M. (2016). Nucleus Accumbens Core and Shell Differentially Encode Reward-Associated Cues after Reinforcer Devaluation. Journal of Neuroscience, 36(4), 1128–1139. 10.1523/JNEUROSCI.2976-15.2016

Willard, F. S., Willard, M. D., Kimple, A. J., Soundararajan, M., Oestreich, E. A., Li, X., Sowa, N. A., Kimple, R. J., Doyle, D. A., Der, C. J., Zylka, M. J., Snider, W. D., & Siderovski, D. P. (2009). Regulator of G-protein signaling 14 (RGS14) is a selective H-Ras effector. PloS One, 4(3), e4884–e4884. 10.1371/journal.pone.0004884

Willis, M. A., & Haines, D. E. (2018). Chapter 31—The Limbic System. In D. E. Haines & G. A. Mihailoff (Eds.), Fundamental Neuroscience for Basic and Clinical Applications (Fifth Edition) (pp. 457–467.e1). Elsevier. 10.1016/B978-0-323-39632-5.00031-1

Xu, J., Kang, N., Jiang, L., Nedergaard, M., & Kang, J. (2005). Activity-dependent long-term potentiation of intrinsic excitability in hippocampal CA1 pyramidal neurons. The Journal of Neuroscience: The Official Journal of the Society for Neuroscience, 25(7), 1750–1760. 10.1523/JNEUROSCI.4217-04.2005

Young, C. B., Reddy, V., & Sonne, J. (2024). Neuroanatomy, Basal Ganglia. In StatPearls. StatPearls Publishing. http://www.ncbi.nlm.nih.gov/books/NBK537141/

Zhang, J., Ding, M., Liu, H., Shi, J., Wang, Z., Wen, P., Zhang, Y., Yan, B., Guo, D., Zhang, X., Tao, R., Yan, Z., Zhang, Y., Liu, Z., Guo, W., & Zhang, S. (2022). Regulator of G-protein signaling 14 protects the liver from ischemia–reperfusion injury by suppressing TGF-β-activated kinase 1 activation. Hepatology (Baltimore, Md.), 75(2), 338–352. 10.1002/hep.32133

